# A multivalent mRNA-LNP therapeutic vaccine with broad cross-genotype immunogenicity elicits clearance of HBV infected hepatocytes

**DOI:** 10.64898/2025.12.03.692117

**Authors:** Daniel Makrinos, Yanbo Sun, Heather Davis, LingZhi Ma, Thu Kim, Emma Dellea, Mia Malone, Edison Ong, Maria Cavallaro, Caroline Atyeo, Phillipa Martin, Cathal Harmon, Ziqiu Wang, Guha Asthagiri Arunkumar, Alec W Freyn, Yen-Ting Lai, Kath Hardcastle, Galit Alter, Andrea Carfi, Anthony DiPiazza, Simone Pecetta

**Affiliations:** Moderna Inc., Cambridge, MA, USA

## Abstract

Chronic hepatitis B remains a major global health challenge, affecting over 254 million individuals and causing over 1 million deaths annually. Despite current antiviral therapies effectively suppressing viral replication, functional cure rates are low due to HBV-induced immune dysfunction and exhaustion. Therefore, new therapeutic approaches to achieve immune control of HBV infection are needed.

Following the systematic evaluation of multiple HBV mRNA antigen designs, we developed mRNA-1965, a trivalent therapeutic mRNA vaccine encoding nanoparticle-displayed PreS1 and PreS2 domains of HBsAg to bypass the immune interference caused by HBV subviral particles, along with mutant forms of HBV Core and Polymerase. In HBV naïve mice, mRNA-1965 immunization induced dose-dependent HBV-neutralizing antibodies and Th1-skewed CD4+ and IFNγ+ CD8+ T cell responses to all three encoded HBV antigens. In non-human primates, mRNA-1965 elicited broad antibody and T cell responses across multiple HBV genotypes. Furthermore, vaccination with mRNA-1965 achieved a strong neutralizing antibody response and complete clearance of serum and liver HBV biomarkers in in an AAV-HBV mouse model with ∼100 IU/mL baseline HBsAg. Notably, combining mRNA-1965 with immune stimulatory co-modalities targeting PD-L1 and OX40 further enhanced therapeutic efficacy in mice with ∼1000 IU/mL baseline HBsAg. Clearance of HBV in AAV-HBV mice was associated with T cell response to mRNA-encoded antigens and with activation and differentiation of Core-specific CD8+ T cells.

These findings support the potential of mRNA-1965 to promote a functional cure for chronic hepatitis B by overcoming immune dysfunction and subsequently enabling robust, functional immunity.

## Introduction

Chronic hepatitis B virus (HBV) infection (CHB) is a serious public health issue, contributing to morbidity and mortality through liver cirrhosis and hepatocellular carcinoma (*1*), as well as increased risk of chronic kidney disease (*2*). Globally, in 2022, there were an estimated 254 million people living with chronic HBV infection, 1.2 million new HBV infections, and approximately 1.1 million deaths attributable to HBV (*3*). Moreover, the annual global deaths from HBV are expected to increase by 39% from 2015 to 2030 (*4*). Prophylactic vaccines are nearly 100% effective in preventing HBV infection, and implementation of vaccination programs over the last 30 years has led to a decline in HBV infections among infants (*5*). However, vaccination coverage remains low in many regions, leaving large portions of the population susceptible to infection through adulthood (*6, 7*).

Elimination of CHB is challenging due to persistent viral replication and a defective HBV specific immune response that results from the combination of high antigen-dose exhaustion and liver-tolerizing pathways (*8, 9*). In particular, HBV subviral particles (SVPs) primarily composed of filaments and spherical aggregates of the Small isoform of Hepatitis B surface antigen (HBsAg) are main contributors of T cell exhaustion (*10*). Published evidence from animal models and subjects that cleared HBV infection, either acute or chronic, suggests that a multifunctional immune response is necessary to achieve cure. In chimpanzees, CD8+ T cells are essential for the control of HBV, driving both cytolytic elimination of HBV and viral clearance via non-cytolytic, IFN-γ-mediated, pathways (*11*). Similarly, in humans, CD8+ T-cell responses against HBV Polymerase, Core, and Surface antigens were identified following clinical recovery and persisted for decades (*12*). Moreover, higher frequencies of functional HBV-specific CD4+ T cells were observed in subjects that achieved functional cure as compared to those who did not (*13*).

Nucleos(t)ide analogues (NUCs) have been widely used to treat chronic HBV infection. Although NUC therapy can profoundly suppress HBV DNA, individuals on NUCs rarely clear HBsAg. Indeed, the annual HBsAg seroclearance rate, a marker associated with HBV “functional cure,” is only 1% (*14*). This leaves patients with a lifelong therapy dependency and limited chances of stopping treatment without risking disease progression, as the risk of hepatocarcinogenesis cannot be eliminated even in patients under strict HBV control (*15*). This is associated with HBV DNA integration, ER stress, and oncogenic HBV proteins in infected hepatocytes, as well as persistent hepatic inflammation, and liver cirrhosis development (*16–21*). Another approved therapeutic option, pegylated interferon (pegIFN), used to stimulate innate immune cells, can be used in combination with NUCs to stimulate the immune system, breaking tolerance and improving rates of functional cure. However, the unfavorable safety profile and the limited success of clearing HBsAg limit its therapeutic use (*22*). Therefore, there is a critical need for new therapeutic strategies to improve upon existing long-term antiviral therapy and the limited effectiveness of pegIFN treatment.

Novel investigational treatments that include entry inhibitors, RNA interference, capsid assembly modulators, and immunomodulatory approaches are being investigated (*23*). Among these, therapeutic vaccination is also being investigated with the aim of subverting virus-induced immune exhaustion and eliminating the viral reservoir. Therapeutic vaccination is expected to work in synergy with direct acting modalities in a sequential “triple therapy combination”, consisting of viral replication and HBV antigen expression suppression followed by immune stimulation (*24*). Indeed, viral-vectored vaccines in sequential combination regimens have recently demonstrated potential for achieving functional HBV cure (*25*). Moreover, combination of immune stimulatory signals improved vaccine immunogenicity, as shown with checkpoint blockade and IFN-a combination therapies (*26, 27*). The former approach draws from the cancer immunotherapy field and has the aim to reinvigorate T cells as well as protect T cells that traffic to the liver (*24, 28, 29*).

Guided by biological and clinical evidence, we developed a therapeutic mRNA-lipid nanoparticle (mRNA-LNP) vaccine for CHB treatment. mRNA-LNP technology is emerging as a potential novel HBV therapeutic vaccine modality. This technology has the advantage of presenting antigens in their native conformation to the immune system, has been shown to elicit strong antibody and cellular immunity, allows for dosing multiple times without the limitation of anti-vector immunity, and enables rapid vaccine optimization. Recent preclinical results demonstrated its potential in murine surrogate chronic HBV models (*30–32*). Our strategy encompassed the systematic testing of multiple rationale HBV antigen designs and their combination in a multivalent mRNA-LNP to obtain a therapeutic vaccine, termed mRNA-1965, that combines synergistic mechanisms to overcome high HBV antigen exhaustion, and we evaluated the potential of co-immunostimulatory signals to potentiate the mRNA vaccine therapeutic efficacy.

## Results

### PreS1 and PreS2-focused mRNA vaccines, as compared to S-HBsAg mRNA or recombinant vaccines, elicit HBV neutralizing antibodies more resistant to HBV SVP interference

HBsAg is a key driver of immunity both in prophylactic and therapeutic HBV vaccines. On the surface of HBV virion, HBsAg is present in three isoforms, Large or L-HBsAg (containing PreS1, PreS2, and Small domains), Medium or M-HBsAg (PreS2 and Small), and Small or S-HBsAg, the latter being the most abundant (*33, 34*). PreS1 contains the human NTCP receptor binding domain and mediates high-affinity interactions that enables HBV entry into the hepatocytes. S-HBsAg contains the “α determinant”, a highly conserved region that facilitate the early low-affinity virion attachment to hepatocytes via heparan sulfate proteoglycans (HSPGs) binding (*35*). Antibodies targeting PreS1 and the “α determinant” block HBV infectivity, the latter being the primary correlate of protection by S-HBsAg-containing prophylactic vaccines (*36–40*).

To determine the immunogenicity of different isoforms of HBsAg encoded with mRNA, we designed four constructs to be expressed as extracellular secreted proteins or membrane-anchored antigens: the transmembrane L-HBsAg antigen, the secreted PreS1 and PreS2 domains of HBsAg displayed on Lumazine Synthase or Ferritin nanoparticles (abbreviated as PreS1S2-LuS and PreS1S2-Fer, respectively), and the secreted S-HBsAg (**Fig. 1A**). These constructs were designed from HBV genotype A HBsAg sequences, to allow comparison with commercial recombinant S-HBsAg Alum-adjuvanted Recombivax vaccine. All constructs elicited strong antibody response in CB6F1 mice immunized twice with the mRNA HBsAg constructs or Recombivax. All PreS1-containing mRNAs elicited comparable, durable PreS1-PreS2-specific antibody IgG titer and, similarly, all S-HBsAg-containing mRNAs elicited durable S-HBsAg-specific antibody IgG titer up to day 226 (**Fig. S1A-B**). A lower Small-specific IgG titer was observed for the L-HBsAg mRNA as compared to the S-HBsAg mRNA, the latter being non-inferior to Recombivax (**Fig. S1B**).

**Figure 1.**
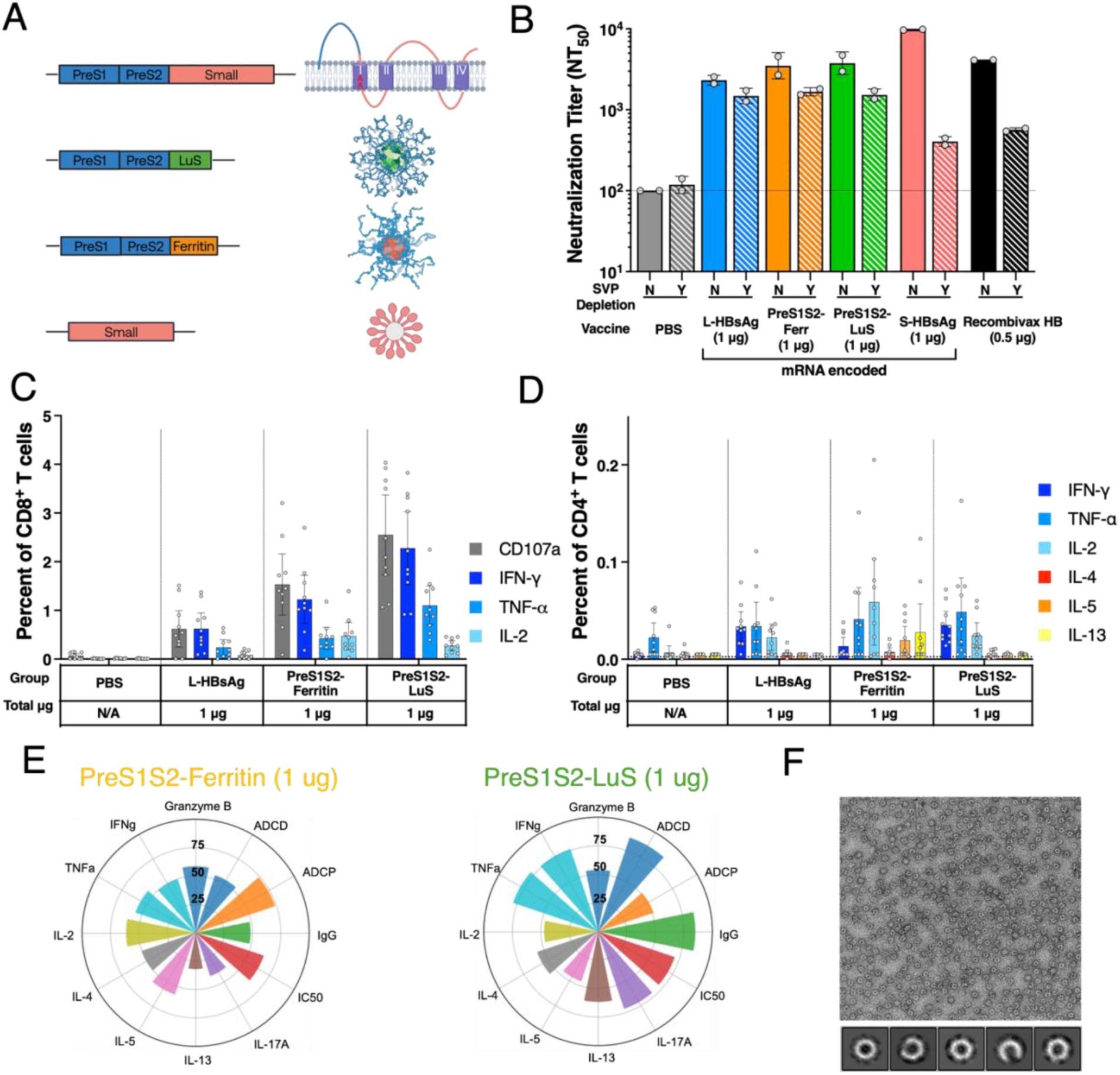
Monovalent HBV HBsAg mRNA vaccines immunogenicity. Female CB6F1 mice received two intramuscular 1 ug doses of HBV mRNA-LNP constructs or 0.5 ug doses of Recombivax or PBS on days 0 and 21. (**A**) Schematic of the mRNA constructs. 2Ala: Alanine substitution; HATM: Influenza HA transmembrane domain. (**B**) Neutralization titer was measured using day 225 serum with or without prior incubation with 33ug/mL of S-HBsAg SVP. (**C-D**) CD8 and CD4 T cell response against PreS1-PreS2 peptides of HBV genotype A was measured by ICS. (**E**) Flower plot summarizing IgG titer, antibody-dependent compliment deposition (ADCD) and phagocytosis (ADCP) measured using day 42 serum. (**F**) Negative stain EM images of PreS1S2-LuS nanoparticles. Plots in panels B-D show group geometric mean with 95% CI for n=10 per group, with assay LLOD indicated by dashed lines.

To determine whether the mRNA-elicited antibodies were able to block HBV infection in the context of CHB, which is characterized by an overabundance of S-HBsAg-rich SVPs (*10*), a live HBV (genotype D) was developed using NTCP-expressing human hepatocyte cell to measure the neutralization potency of mice sera in presence of physiologically relevant concentration of SVPs. Only genotype D HBV virus was used in this assay, as live virus stocks of other genotypes were not available. A marked 24.1- and 7.3-fold reduction in neutralization titer was observed for sera of mice immunized with S-HBsAg mRNA or Recombivax, respectively (**Fig. 1B**). This evidence is in line with *in vitro* HBV SVPs-driven reduction of human S-HBsAg vaccinated serum neutralization (*41*). Conversely, a 1.6-2.5-fold reduction of neutralization titer was observed for sera of mice immunized with PreS1-PreS2-containing mRNA vaccines, suggesting efficient viral blocking despite the presence of SVPs (**Fig. 1B**). Therefore, we decided to focus on PreS1-containing mRNA construct for further vaccine development.

We then evaluated the magnitude of HBV-specific T cell response induced in mice immunized twice with the mRNA vaccines via intracellular cytokine staining upon in vitro stimulation of splenocytes collected at Day 225 in presence of HBV antigen peptides. PreS1S2-LuS induced a superior magnitude of CD8+ T cells, including IFNy and TNFa, with 2% mouse splenocytes specific for PreS1-PreS2 peptides, as compared to L-HBsAg and PreS1S2-Fer mRNAs, 0.5% and 1%, respectively (**Fig. 1C**). The PreS1-PreS2-specific CD4+ T cell response was comparable for these antigens, 0.02% IFN-y on average, primarily Th1-skewed, though a Th2 response was also observed for the PreS1S2-Fer mRNA (**Fig. 1D**). Of note, the L-HBsAg and S-HBsAg also generated strong S-HBsAg-specific CD8+ T cell responses, which instead were absent for Recombivax (**Fig. S1C)**. The S-HBsAg-specific CD4+ response was low and comparable among antigens, though a difference in phenotype was observed, with mRNA immunization inducing a Th1-skewed response while a Th1-Th2 response was induced by Recombivax (**Fig. S1D)**.

The combination of both humoral and cellular immune data reveals that all mRNA-encoded constructs induced a multi-functional immunity, with antibodies capable of neutralize HBV, and with robust IFNy+ TNFa+ CD8+ and Th1-skewed CD4+ T cell response which are recognized as critical for HBV elimination. The PreS1-PreS2-contaning mRNAs were surprisingly potent immunogens and, to further evaluate their immunogenicity, we measured antibody-dependent complement deposition (ADCD) and cellular phagocytosis (ADCP) using reporter assays. Antibodies elicited by PreS1S2-LuS and PreS1S2-Ferritin mRNAs provided the strong ADCD and ADCP scores against PreS1-PreS2-cells expressing the target antigen (**Fig. S1E-F**). Overall, the PreS1S2-LuS design surpassed the PreS1S2-Fer on the account of multiple immunological readouts. It is well validated that antigen presentation on a particulate antigen has been reported to enhance immunogenicity (*42*). To confirm that the PreS1S2-LuS self-assembles into a 60-mer nanosphere, we purified the PreS1S2-LuS from the supernatant of mRNA transfected Expi293 cells and negative staining microscopy analysis revealed well-folded, correctly sized nanoparticles (**Fig 1F**). In consideration of its immunogenicity, and the evidence that PreS1-PreS2-targeting antibodies are likely more effective than Small-targeting antibodies in the context of CHB, we therefore elected the PreS1S2-LuS mRNA construct for further vaccine development.

### HBV Core and Polymerase mRNAs elicit strong IFNy+ CD8+ and Th1-skewed CD4+ T cell response

As demonstrated in animal models and human samples, T cell response is critical for the elimination of chronic HBV infection. To complement the strong T cell immunity provided by the PreS1S2-LuS mRNA, we designed mRNA-encoded HBV Core and Polymerase constructs (**Fig 2A-B**).

**Figure 2.**
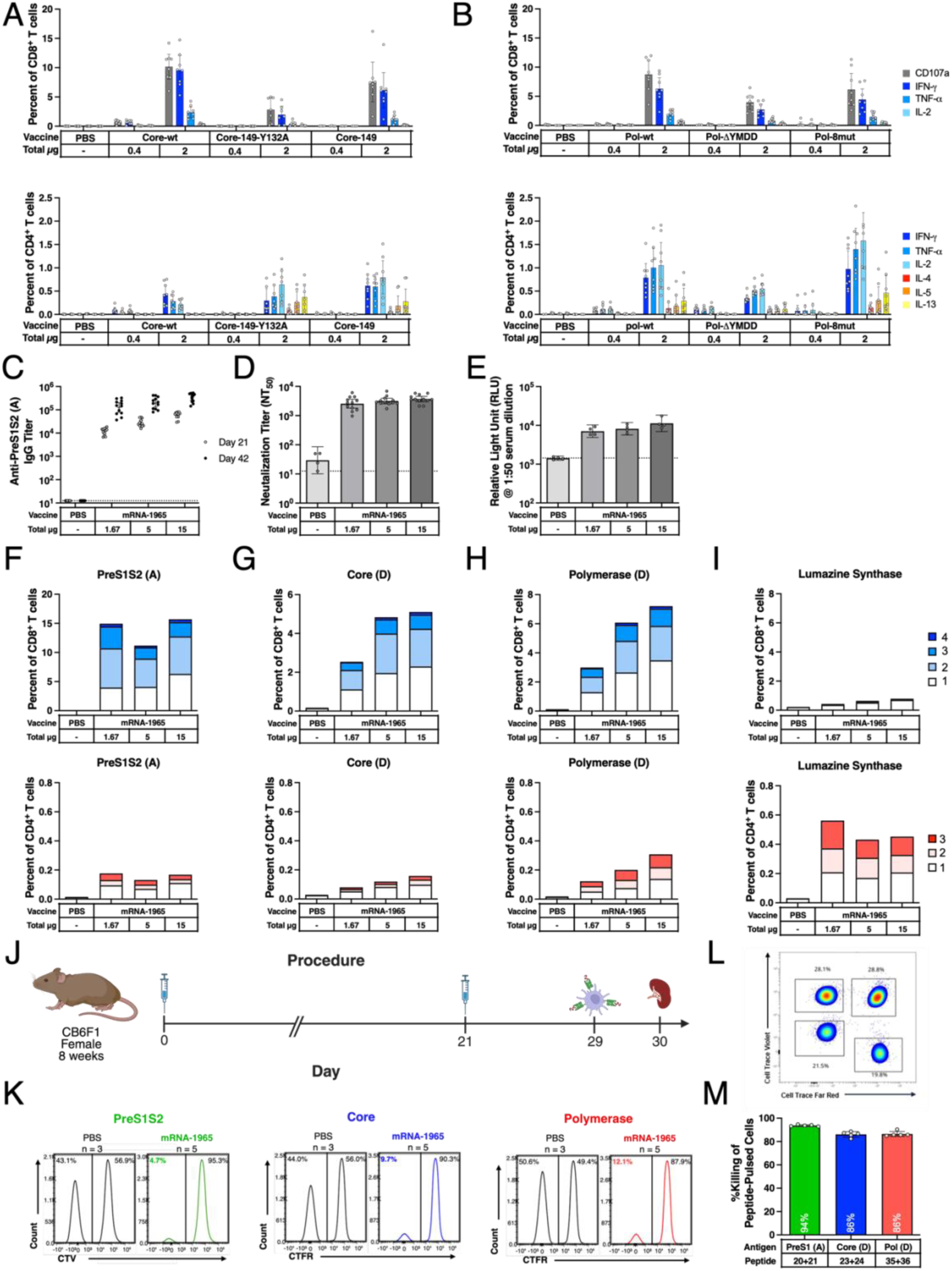
Monovalent and Multivalent HBV mRNA vaccines immunogenicity. (**A-B**) Mice received two doses of mRNA constructs of HBV Core (**A**) or Polymerase (**B**) or PBS on days 0 and 28. CD8 (**Top**) and CD4 (**Bottom**) T cell response was measured by ICS. (**C-I**) Mice received trivalent mRNA-1965 or PBS on days 0 and 28. (**C**) Anti-PreS1S2 (genotype A) IgG binding titers. (**D-E**) Day 42 serum neutralization titer (**D**) and ADCC (**E**). (**F-I**) CD8^+^ (**Top**) and CD4^+^ (**Bottom**) T cell response against PreS1S2 (genotype A) (**F**), Core (**G**) and Polymerase (**H**) (genotype D), and Lumazine synthase (**I**) were measured by ICS assay (Day 42 splenocytes). Colors denotes the proportion of cells expressing respective number of CD8+ (CD107a, IFN-γ, TNF-α, IL-2) or Th1 CD4+ (IFN-γ, TNF-α, IL-2) T cell markers. (**J-M**) Mice received mRNA-1965 (15 ug) or PBS on days 0 and 21, followed by adoptive transfer (day 28) mosaic-labeled splenocytes pulsed with dominant epitopes of preS1S2, Core, Polymerase, or COVID Spike. (**J**) Study schematic. (**K**) Representative flow data showing mosaic labeling. (**L-M**) Representative flow histogram (**L**) and quantification (**M**) of antigen-specific killing. Plots show group geometric mean with 95% CI with assay LLOD indicated by dashed lines. n=8 **(A-B)**, n=12 **(C-I)**, n=5 **(J-M)** per group.

We first evaluated the magnitude of HBV-specific T cell response induced in mice immunized twice with three monovalent HBV Core and three monovalent Polymerase constructs, which are designed to be expressed as cytosolic proteins. The three HBV Core mRNAs encoded for either wt Core genotype D sequence, a truncated Core in amino acid position 149 to eliminate the DNA-binding motif while still allowing capsid assembly (*43*), and a truncated Core with a Y132A mutation that abolishes capsid assembly (*44*). All three designs induced robust antigen-specific Th1-skewed CD4+ and high IFNy+ CD8+ T cell responses, the latter representing 5-10% HBV-specific among all T cells in CB6F1 immunized mice splenocytes sampled at day 35 (**Fig. 2A**). HBV Core-149 mRNA resulted in a balanced CD4+/CD8+ T cell immunogenicity and selected for further development, while the Core-wt and the Core-149-Y132A mRNAs demonstrated a CD8-dominant or CD4-dominant T cell immunogenicity, respectively. Comparably, the three HBV Polymerase mRNA designs, a wt Polymerase genotype D sequence, a Polymerase mutant with 8 amino acid substitutions that abolish its catalytic activity (*45*), and a second Polymerase mutant with a ΔYMDD deletion in the reverse transcriptase domain (*46*), induced robust antigen-specific Th1-dominant CD4+ and IFNy+ CD8+ T cell response following mouse immunization (**Fig. 2B**). Given its strong and well-balanced CD8 and CD4 T cell response, the inactive Pol-8mut was preferred for further development. As an additional strategy, we also expressed a PreS1S2-Core nanoparticle design that displayed PreS1 and PreS2 on the HBV Core antigen, either wild type or 149 mutant, with the aim to combine these two antigens in one construct. The HBV Core has been extensively used in the literature to scaffold antigens, and it has higher a valency (180-240-mers) as compared to both ferritin (24-mer) and Lumazine synthase (60-mer) (*47, 48*). PreS1S2-LuS, PreS1S2-Core-wt, and PreS1S2-Core149 induced robust neutralizing antibodies and PreS1S2-specific CD8^+^ and CD4^+^ T cell responses. However, the Core-specific T cell response was reduced as compared to HBV Core mRNA alone, therefore we opted to keep the PreS1S2 and Core antigens separated (**Fig S2**).

These results demonstrate that all mRNA-encoded HBV cytosolic antigens are robustly immunogenic eliciting strong HBV-specific functional T cell immunity. To further validate these mRNAs, we combined the three selected antigens, PreS1S2-LuS, Core-149, and Pol-8mut, in a trivalent mRNA-LNP formulation named mRNA-1965.

### The trivalent mRNA vaccine, mRNA-1965, elicits multifunctional dose-dependent immunity in mice against all encoded HBV antigens

To evaluate the immunogenicity of mRNA-1965, we analyzed HBV-specific serum antibody levels and T cell response in CB6F1 mice immunized twice with the trivalent mRNA-LNP formulation. Immunization with mRNA-1965 elicited serum IgG binding antibodies to the PreS1 and PreS2 domains of HBsAg, with a dose-dependent antibody titer (**Fig. 2C**). These antibodies exhibited HBV neutralization at all tested doses (**Fig. 2D**) which, as assessed in a separate study, were non-inferior to monovalent PreS1S2-LuS mRNA immunization (**Fig. S3A-C**). Similarly, serum antibodies elicited by mRNA-1965 exhibited PreS1-specific ADCC Fc effector function at all tested doses (**Fig. 2E**). In line with monovalent mRNAs results, mRNA-1965 induced strong CD8+ and CD4+ T-cell responses specific for HBV PreS1S2, HBV Core and HBV Polymerase peptides in mouse splenocytes sampled at 14 days post-second dose (**Fig. 2F-H**). A CD4+ but not CD8+ T cell response was observed for Lumazine Synthase peptides (**Fig. 2I**). In a separate study, we observed that the combination of mRNAs resulted in a reduced magnitude of Core- and Polymerase-specific T cell response when compared to single mRNAs, though this interference was partially restored with at high dose (**Fig. S3D-E**).

To validate the evidence that mRNA-1965-induced CD8+ T-cells are indeed able to specifically eliminate HBV-presenting cells, we immunized CB6F1 mice twice with mRNA-1965 and adoptively transferred labeled antigen presenting cells (APCs) pulsed with HBV peptides 7 days post-2^nd^ dose (**Fig 2J-K**). These peptides were previously identified as highly immunogenic CD8 IFN-y inducing epitopes in the CB6F1 mouse strain (**Fig S4**). A near complete elimination of HBV-peptide pulsed APCs was observed 16 hours following adoptive transfer, with 94% PreS1, 86% Core, and 86% target cell reduction, respectively, denoting highly functional cytolytic activity elicited by mRNA-1965 (**Fig 2L-M**).

These results demonstrate that mRNA-1965 elicits a multi-functional immunity characterized by HBV neutralizing antibodies, PreS1-PreS2-targeting antibodies that are capable of inducing PreS1-specific ADCC, and HBV PreS1-PreS2, Core, Polymerase specific CD4+ and CD8+ T-cells, the latter with strong cytolytic activity. These mechanisms could result in control of HBV infection via cell-to-cell transmission blockage and antibody- and T cell-mediated HBV-infected hepatocyte elimination.

### mRNA-1965 elicits cross-genotype antibodies and T cells in non-human primates

HBV is classified into ten distinct genotypes (A–J), each with a unique geographic distribution and potential implications for disease progression and treatment response (*49*). To evaluate the breadth of immune response to mRNA-1965, which encodes genotype A PreS1-PreS2 and genotype D Core and Polymerase sequences, across this genetic diversity, cynomolgus macaques of Cambodian origin received three mRNA-1965 doses, and their humoral and cellular immune response were accessed (**Fig. 3A**).

**Figure 3.**
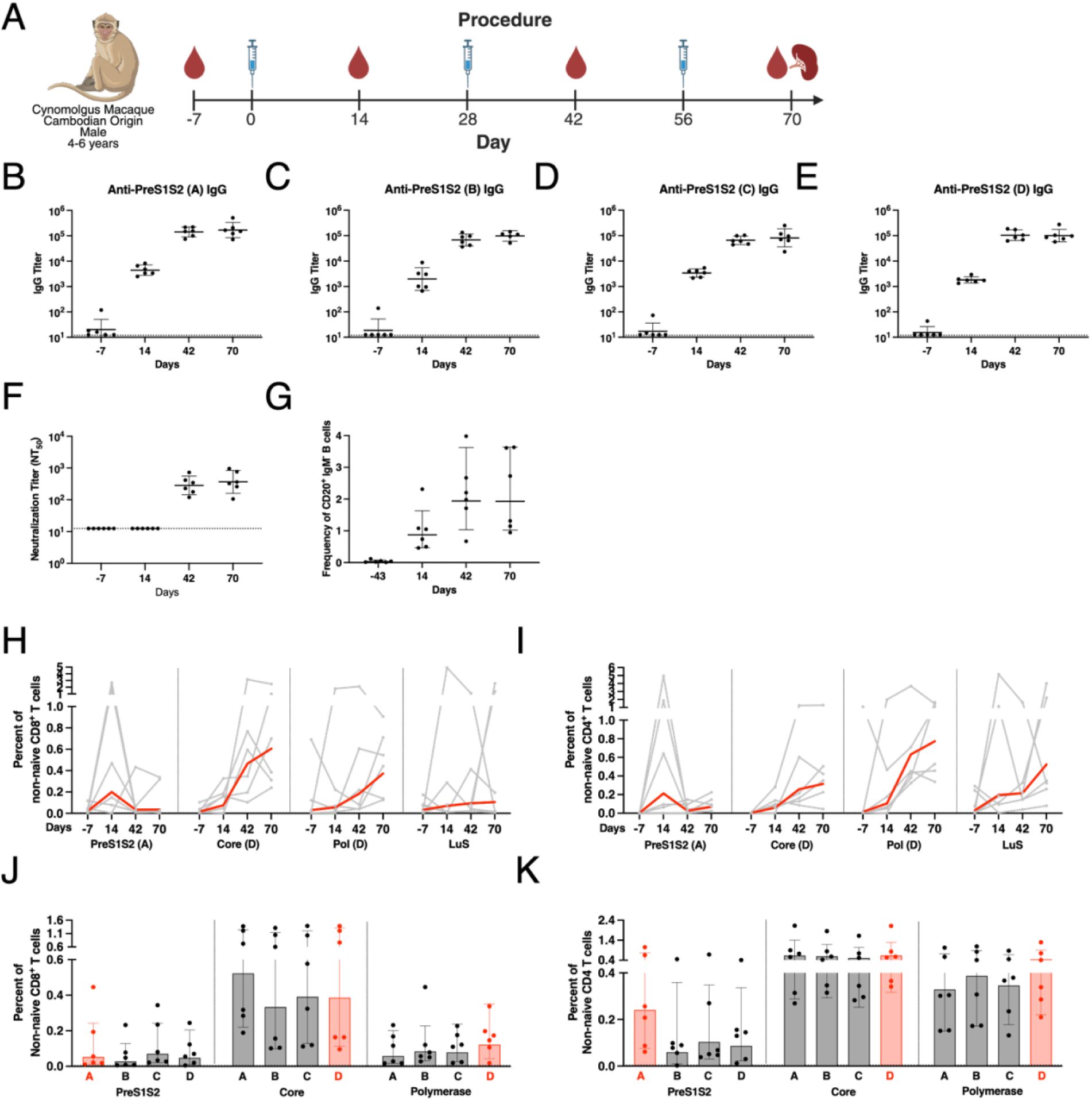
mRNA-1965 immunogenicity in NHPs. Cynomolgus Macaques received three doses of mRNA-1965 (150ug). (**A**) Study schematic. (**B-E**) IgG binding titers against PreS1S2 of HBV genotypes A (**B**), B (**C**), C (**D**), and D (**E**). (**F**) Neutralization titers. (**G**) PreS1S2-specific class-switched B cells. (**H-I**) "Any CTL" CD8+ (**H**) and "Any Th1" CD4+ (**I**) T cell responses to PreS1S2 (genotype A), Core (genotype D), Polymerase (genotype D), and Lumazine Synthase in PBMC samples. (**J-K**) "Any CTL" CD8+ (**J**) and "Any Th1" CD4+ (**K**) T cell responses to PreS1S2, Core, and Polymerase (genotypes A–D) in splenocyte samples. Red bars indicate genotypes matching those encoded in mRNA-1965. (**B-K**) Each dot or grey line in the plot represents an individual macaque, with the bold or red line or bar height and error margins represent the geometric mean with 95% confidence intervals (CI). n=6. Assay LLODs are indicated by the dashed lines. (**H, J**) "Any CTL" CD8+ T cell responses are defined by positivity of at least one following markers (CD107a, IFN-γ, TNF-α, IL-2). (**I, K**) "Any Th1" CD4+ T cell responses are defined by positivity of at least one following markers (IFN-γ, TNF-α, IL-2, CD40L). (**J-K**) Data was analyzed by two-way ANOVA with Tukey’s correction.

Immunization of non-human primates (NHPs) with mRNA did not result in significant body temperature or weight changes, suggesting that the vaccine was well tolerated (**Fig. S5A-B**). Serum IgG antibodies targeting HBV PreS1-PreS2 were elicited following mRNA-1965 immunization, with cross-reactivity across HBV genotypes A–D (**Fig. 3B-E**), the most common genotypes in the North American, European, and Asia-Pacific regions (*50*). Antibody titers and kinetics were comparable among genotypes, with peak responses observed after the first booster dose (**Fig. 3B-E**). Neutralization titer against live HBV virus, genotype D, was observed following the booster doses (**Fig. 3F**). These antibody responses were accompanied by a proportional increase of memory B cell response denoting robust germinal center reaction (**Fig. 3G**). Peripheral antigen-specific T cell response to vaccine-matched PreS1-PreS2 peptides was generally low and peaked following prime dose while a dose-dependent T cell response to vaccine-matched Core and Polymerase peptides was observed (**Fig. 3H-I**). Despite a lower PreS1S2-LuS T cell immunogenicity in NHPs as compared to CB6F1 mice, the CD4+ T cell responses were characterized by a Th1 phenotype and the CD8+ T cell response were characterized by a cytotoxic phenotype, in accordance with murine immunogenicity data. Nonetheless, when splenocyte T cells were analyzed at terminal sampling, immunity to all antigens was detected, which was comparably cross-reactive against genotype A-D peptides (**Fig. 3J-K**). Interestingly, despite a lower magnitude of T cell response to PreS1-PreS2, a strong T cell immunity to Lumazine synthase was detected (**Fig. 3H-I**). This evidence, together with strong antibody and memory B cell response, suggests that Lumazine synthase-specific T cells may participate in B cell cross-priming – a phenomenon that has been well-documented in the literature (*51, 52*).

Taken together, these results demonstrate that mRNA-1965 is safe and immunogenic in NHPs in a three-dose regimen, with induction of cross-genotype reactive antibodies and T cells. Of note, a CD8+ T cell epitope prediction analysis suggest a high degree of HBV genotype cross-reactivity in humans (**Fig. S5C**). Interestingly, a difference in PreS1-S2 T cell immunogenicity was observed between species, though the inclusion of a non-HBV antigen may help in driving a robust and functional neutralizing antibody response, which could be an advantage in chronic hepatitis B.

### mRNA-1965 monotherapy clears serum and hepatic HBV biomarkers in AAV-HBV mice with low baseline level of viremia and antigenemia

To determine the immune response to mRNA-1965 in the context of chronic HBV infection, we developed an adenovirus associated surrogate chronic hepatitis B mouse model (referred as AAV-HBV) to evaluate the therapeutic efficacy of mRNA-1965. A preliminary analysis of the level of serum HBsAg in mice injected with AAV-HBV (genotype D) virus revealed a low antigenemia in CB6F1 mice as compared to C57B6 male mice (**Fig. S6**). Therefore, to align with the AAV-HBV body of literature, we opted for using C57BL/6 mice for the following experiments.

To investigate the role of increasing HBV burden to mRNA immunogenicity in this model, we injected ascending doses of AAV-HBV, and obtained a laddered serum HBV biomarker levels in the AAV-HBV mice, ranging from 10 to 10,000 IU/mL serum HBsAg, 10-1000 IU/mL serum HBeAg, and undetectable to 10^6^ HBV DNA copies/mL (**Fig. 4A-D, S7A-C**). When the AAV-HBV mice were later immunized with four doses of mRNA-1965 a complete elimination of serum HBV markers was observed in mice with low antigenemia (<100 IU/mL HBsAg, <100 IU/mL HBeAg) and viremia (undetectable HBV DNA), which occurred after two or four mRNA-1965 doses (**Fig. 4B-D**). However, mRNA-1965 alone elicited only a transient reduction of serum HBV biomarkers in mice with high antigenemia (>1000 IU/mL HBsAg, >100 IU/mL HBeAg) and viremia (>10,000 IU/mL), with viral rebound observed by the end of treatment (**Fig. 4B-D**).

**Figure 4.**
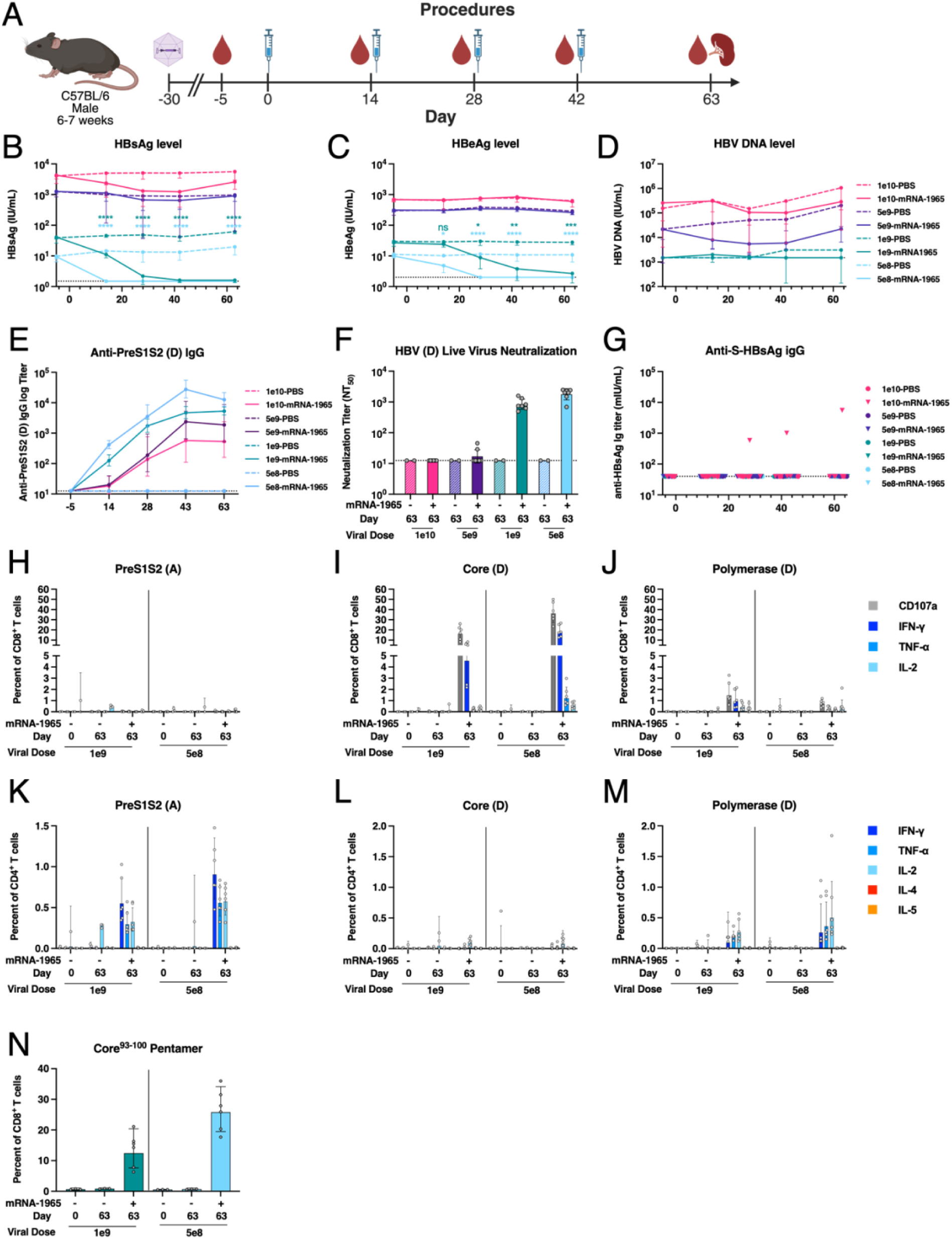
Therapeutic efficacy of mRNA-1965 in AAV-HBV mouse model. Mice received 1e10, 5e9, 1e9, or 5e8 vector genomes of recombinant AAV-HBV on Day -30, followed by four intramuscular doses of mRNA-1965 (15ug) or PBS. (**A**) Study schematic. (**B-D**) Serum HBsAg (**B**), HBeAg (**C**), and HBV DNA (**D**) levels. (**E**) Anti-PreS1S2 (Genotype D) IgG binding titers. (**F**) Neutralization titers. (**G**) Anti-S-HBsAg IgG binding titers. (**H-M**) CD8+ (**H-J**) and CD4+ (**K-M**) T cell responses to PreS1S2 (genotype A) (**H, K**), Core (genotype D) (**I, L**), Polymerase (genotype D) (**J, M**) in splenocytes. (**N**) Core^93-100^ pentamer+ CD8+ T cell responses in splenocytes. (**B-N**) n= 7 for mRNA-1965 treated groups, and n=3 for PBS treated groups. Dashed lines denote assay LLODs. (**B-D**) Dot and error margins represent geometric mean with 95% confidence intervals (CI). Statistical analysis was done with one sample t-test with Benjamini-Hochberg correction. * indicates P-value of 0.01-0.05, ** indicates P-value of 0.001-0.01, *** indicates P-value of 0.0001-0.001, **** indicates P-value smaller than 0.0001. (**E-N**) Each dot represents an individual mouse, with the bold line or bar height and error margins represent the geometric mean with 95% confidence intervals (CI).

We next assessed the HBV-specific humoral immune response. Notably, all mRNA-1965–treated mice exhibited a robust IgG response against the PreS1-PreS2 domains of HBsAg derived from genotype D, which matches the genotype of the AAV-encoded HBV but differs from the genotype A antigen encoded by mRNA-1965 (**Fig. 4E**). Furthermore, we detected strong serum neutralizing activity in mice that achieved clearance of HBV biomarkers following mRNA-1965 treatment (**Fig. 4F**). In contrast, S-HBsAg (pan-genotype)-specific IgG was undetectable in all groups, except for a single mRNA-1965–treated mouse in the high AAV-HBV dose cohort (**Fig. 4G**), indicating that seroclearance of HBsAg occurred independently of an anti-S-HBsAg antibody response.

Given the critical role of cellular immune responses in the clearance of chronic HBV infection, we next characterized HBV-specific CD4+ and CD8+ T cell responses. In mice that cleared HBV biomarkers following mRNA-1965 treatment, we observed a robust CD8+ T cell response targeting HBV Core and Polymerase antigens, with a moderate response to PreS1-PreS2 (**Fig. 4H-J**). Conversely, CD4+ T cells exhibited strong reactivity against PreS1-PreS2 and Polymerase, and to a lesser extent, Core (**Fig. 4K-M**).

It is worth noting that we observed substantially lower CD8+ T cells response in the seroconverted AAV-HBV as well as the vaccinated donor mice used for adoptive transfer than in naïve CB6F1 mice. To further investigate difference in the landscape of cellular immune response to mRNA-1965, we conducted a head-to-head immunogenicity comparison of mRNA-1965 in naïve CB6F1 and C57BL/6 mice. In CB6F1 mice, mRNA-1965 induced strong CD8+ T cell and minimal CD4 T cell response to PreS1S2, moderate CD8+ T cell and minimal CD4+ T cell response to Core, and moderate CD8+ T cell and moderate CD4+ T cell response to Polymerase. In contrast, C57BL/6 mice exhibited a distinct immunogenicity profile, with minimal CD8+ T cell and robust CD4+ T cell response to PreS1S2, robust CD8+ T cell and moderate CD4+ T cell response to Core, and minimal CD8+ T cell and robust CD4+ T cell response to Polymerase (**Fig. S4D**).

To further assess the functionality of vaccine-induced CD8+ T cells, we performed MHC class I pentamer staining targeting a dominant epitope within the Core antigen (**Fig. S4A**). The frequency of antigen-specific CD8+ T cells measured by pentamer staining was comparable to that determined by ICS (**Fig. 4N**), suggesting that the majority of vaccine-induced CD8+ T cells retained functionality in mice that achieved HBV biomarker clearance following mRNA-1965 treatment.

### mRNA-1965 efficacy in high viremic AAV-HBV mice is potentiated by checkpoint blockade co-therapy

Given that mRNA-1965 monotherapy elicited only a transient reduction in serum HBV biomarkers in mice with high baseline antigenemia and viremia, with viral rebound observed following treatment cessation, we next investigated co-modalities to enhance the therapeutic efficacy of mRNA-1965 when administered in combination. In the same study, an additional group of mice at high AAV-HBV dose level received a five doses treatment course with an antibody cocktail containing an OX40 agonist IgG and a PD-L1 antagonist IgG, both targets shown to enhance T cell responses in CHB (*53–56*), administered every three days starting one week after completion of the four-dose mRNA-1965 regimen (**Fig. 5A**).

**Figure 5.**
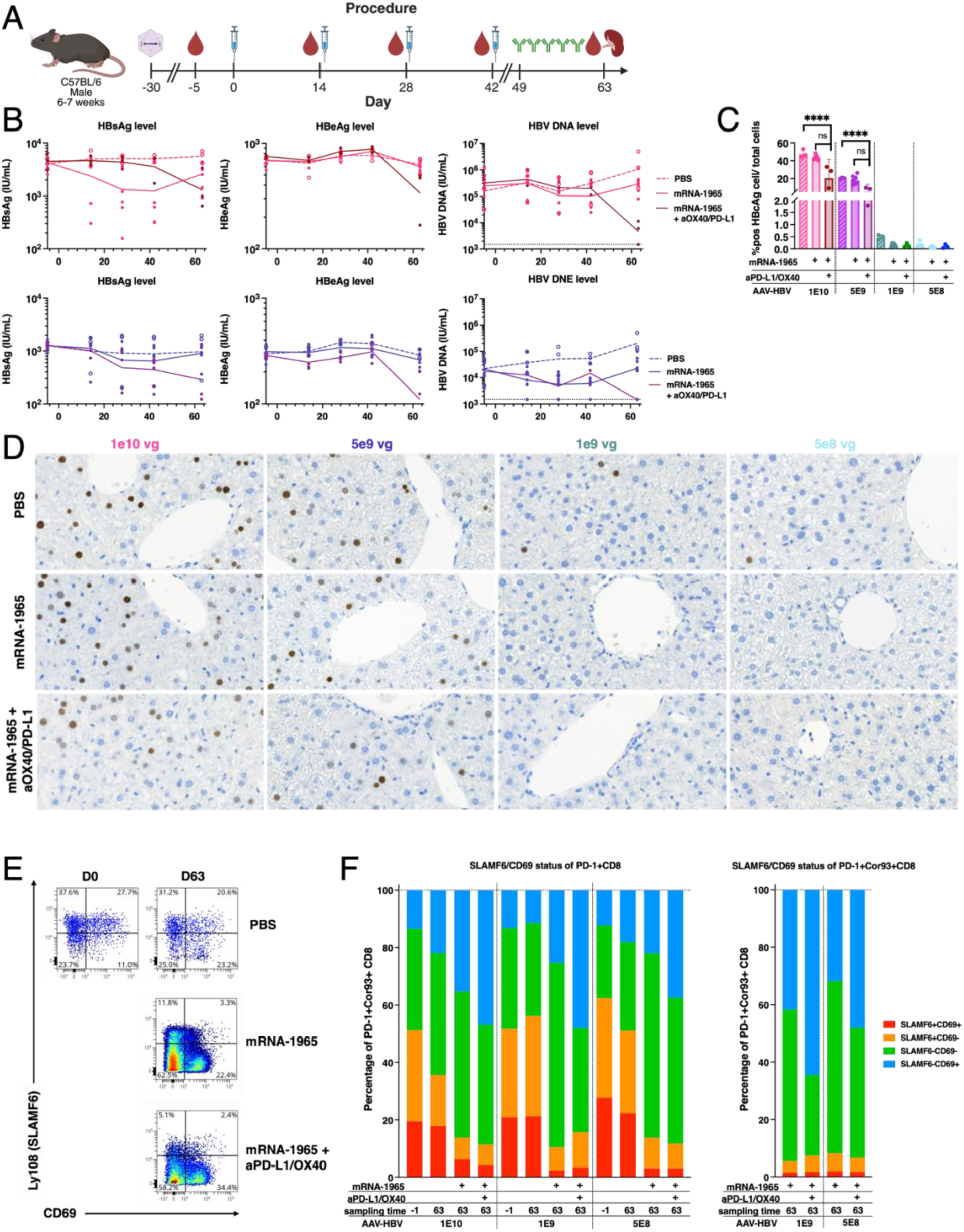
AAV-HBV T cell phenotyping following mRNA-1965 treatment. Male C57BL/6 mice, 6-7 weeks old, were injected through the tail vein with vector genomes of recombinant AAV-HBV on Day -30 as described in Fig 4. In addition to four intramuscular doses of mRNA-1965, a group of mice also received five intraperitoneal doses of a combination of two monoclonal antibodies, an OX40 agonist and a PD-L1 antagonist, administered every 3 days starting on Day 49. Serum samples were collected on Days -5, 14, 28, 42, 63, and splenocyte samples were collected on Day 63. (**A**) Study schematic. (**B**) Serum HBsAg (**Left**), HBeAg (**Middle**), and HBV DNA (**Right**) of groups of mice inoculated with 1e10 vg (**Top**) or 5e9 vg (**Bottom**) of AAV-HBV. Bold line indicates group geometric mean. (**C)** Quantification of percentage HBcAg positive cells within all nucleated cells. (**D**) Representative IHC images of HBcAg staining in FFPE liver samples. Bar height and error margins indicate group geometric mean and 95% CI. (**E**) Representative flow plots for exhaustion status across time points and treatment. (**F**) Quantification of exhaustion status of PD-1+ (**Left**), and Core Core^93-100^ pentamer+ (**Right**) CD8+ T cells across time points and treatment.

At both AAV-HBV inoculation dose, administration of the antibody cocktail led to an approximate 1-log reduction in serum HBsAg levels and a 60–70% reduction in HBV core-positive hepatocytes relative to mRNA-1965 alone (**Fig. 5B-D**). This evidence suggests that checkpoint blockade therapy is effective at potentiating mRNA vaccine-induced immunity, and enhance therapeutic efficacy.

### T cell effector phenotype differentiation correlates with mRNA-1965-induced HBV clearance

It is well established that although the differentiation trajectory of exhausted CD8^+^ T cells, due to chronic viral infection or cancer, differs from that of conventional effector and memory CD8+ T cells, these cells nonetheless undergo a defined path of differentiation upon productive activation (*57*). Distinct stages within this trajectory can be delineated by characteristic cell surface markers. SLAMF6^+^ CD8^+^ T cells, also known as Ly108, CD69+ and CD69-populations are referred to as Tex^prog1^ and Tex^prog2^, respectively. These two subsets are less differentiated and possess higher proliferative capacity when compared to the two SLAMF6-populations. Conversely, the SLAMF6-CD69+ population, referred to as the Tex^term^, is the most differentiated population amongst the four and possesses lowest proliferative capacity. Hence, we sought to characterize the differentiation status of HBV-specific CD8⁺ T cells before and after monotherapy or combination therapy, to identify phenotypic signatures associated with clearance of HBV biomarkers.

Due to the extremely low frequency of Cor93-pentamer⁺ CD8⁺ T cells in pre-treatment samples collected on Day 30 post AAV-HBV inoculation, prior to the first dose of mRNA-1965, and in PBS-treated controls, we instead analyzed PD-1⁺ CD8⁺ T cells as a proxy for HBV-specific population. Using SLAMF6 and CD69 co-expression to infer differentiation status, we observed a similar distribution of the four defined differentiation subsets in both pre-treatment and PBS-treated mice across AAV-HBV dose groups, excluding the 5e9 group due to low yield of splenocytes (**Fig. 5E-F**).

Notably, mRNA-1965 monotherapy resulted in a reduction in the proportion of progenitor exhausted CD8⁺ T cells (SLAMF6⁺ CD69⁺/⁻), suggesting activation and progression along the differentiation trajectory (**Fig. 5E-F**). Furthermore, sequential treatment with the antibody cocktail led to a higher proportion of more terminally differentiated/exhausted CD8⁺ T cells (SLAMF6⁻ CD69⁺) compared to those receiving mRNA-1965 alone in mice with high baseline antigenemia and viremia (1e10 AAV group) (**Fig. 5E-F**). Therefore, mRNA-1965 alone or with combination therapy induced activation and differentiation of phenotypically exhausted Core-specific CD8+ T cell, which is expected to participate in HBV clearance in the context of chronic infection.

## Discussion

HBV vaccines have had a significant impact in preventing liver disease, however, despite showing promising efficacy in preclinical models, they had limited success in treating CHB. Therapeutic use of alum adjuvanted recombinant HBsAg vaccines with reinforced 6-dose immunization schedules elicited HBV serum DNA elimination in a subset of CHB subjects but did not achieve serum HBsAg clearance (*58, 59*). More recently, immunization of NUC controlled CHB subjects with BRII-179, composed of adjuvanted recombinant PreS1, PreS2, and S-HBsAg, achieved elicitation of HBV-specific IFN-y producing T cells in the majority of patients, though limited anti-HBV antibody response and no reduction of serum HBsAg were observed (*27*). Similarly, VTP-300, a chimpanzee adenovirus/MVA-vectored vaccine encoding for the HBV envelope, Core and Polymerase, elicited CD4+ and CD8+ antigen-specific responses which correlated with HBsAg reduction, though most participants in the vaccine-only group did not experience HBsAg change (*26*). Taken together, this evidence underscores the challenge of overcoming immune tolerance caused by chronic exposure to HBV with therapeutic vaccination.

A significant barrier to HBsAg-containing HBV vaccines is the limited antibody and T cell immune response to HBsAg due to formation of antibody-HBsAg complexes and high antigen load-driven immune exhaustion, respectively. To address this challenge, we pursued a strategy that uses mRNA technology to immuno-focus the immune response on the PreS1 and PreS2 domains of HBsAg and elicit T cell immunity to multiple key HBV antigens.

We demonstrated that mRNA can induce strong functional immunity against three key HBV antigens, an approach that also other therapeutic vaccine have employed though with specific advantages offered by the mRNA technology. First, we demonstrated that mRNA can encode designer particles to avoid SVP immune interference. Second, the mRNA encoded HBV antigens elicited robust and broadly reactive CD8 T cell responses across species, which are critical for HBV control, as compared with limited CD8 immunity of recombinant protein with adjuvant vaccines. Third, we demonstrated that the mRNA vaccine can be dosed multiple times and that it can be combined with checkpoint blockade stimuli to boost immunogenicity, an approach that may be necessary to achieve sustained viral control in the context of CHB. Notably, combining mRNA therapeutic vaccine for CHB with ICB could work synergistically to enhance the therapeutic efficacy even further. Recent report has suggested that mRNA-LNP vaccination can sensitize tumors to immune checkpoint blockade therapy, both in clinical cohorts and in pre-clinical mouse models, by increasing in Type-I IFN levels (*60*). This observation further underlines the advantage of mRNA-LNP as a therapeutic vaccine platform for CHB.

HBV infects human hepatocytes via PreS1 binding to the human NTCP receptor. PreS1 is less abundantly expressed on subviral particle (SVP) decoys but highly enriched on infectious Dane particles (*61*), making it an attractive target for virus neutralizing antibodies for CHB treatment and offering a functional advantage over conventional anti-S-HBsAg approaches. PreS1-targeting antibodies, such as 2H5-A14, have been reported to block HBV entry by interfering with the virus-receptor interaction and to decrease the number of HBV-carrying hepatocytes in treated HBV carrying mice. The mechanism of PreS1 antibody-mediated HBV clearance was demonstrated to involve ADCC of liver cells, suggesting that immune-mediated clearance can be achieved even in absence of S-HBsAg antibodies. In other preclinical studies, a PreS1-Ferritin nanoparticle vaccine has been shown to elicit robust PreS1-targeting antibodies and to reduce the number of HBV Core-positive hepatocytes as well as total HBV DNA in the liver, despite the modest impact on serum HBsAg level. The effect was potentiated by suppressing viral antigen expression with siRNA (*62*). The advantage of PreS1-targeting strategy is not limited to preclinical models. Libevitug (HH-003), a PreS1-specific monoclonal antibody, has been approved in China for the treatment of HDV infection, a virus that hijacks HBV Envelope protein to enter hepatocytes, further validating the therapeutic potential of this approach. In addition to PreS1 domain, we included PreS2 domain as additional target for neutralizing antibodies (*63–65*). By expressing PreS1-PreS2 on a nanoparticle scaffold, we achieved robust HBV neutralizing antibody response in presence of SVP both in vitro and in vivo, in absence of S-HBsAg immunity. Our research demonstrates that mRNA can elicit immunity against specific subdomains of HBsAg, either as monomer, full-length membrane-bound antigen, and self-assembly nanoparticles, with the latter inducing a surprising 70-fold enhancement of PreS1-specific antibody response. The use of a non-HBV scaffold as carrier for PreS1 has also other potential advantages in CHB, as described below.

Given there are 10 genotypes of HBV with different geographical distributions, it is crucial to address the cross-genotype reactivity of a therapeutic vaccine to ensure broad efficacy across diverse patient populations. While HBV genotypes B and C are more commonly used in other therapeutic vaccines due to the high proportion of patients with HBV B and C infection, our in-silico epitope analysis suggests that HBV antigens derived from genotypes A through D are likely comparable in immunogenic potential. Indeed, in vivo, we have observed that mRNA-1965 elicited robust and cross-genotype reactive B and T cell response to encoded antigens in Cynomolgus macaque, a model species that exhibits greater HLA complexity than mice and closer immunological similarity to humans (*66*). Nonhuman primates share key features of human immune system, including TCR repertoires, antigen presentation pathways, cytokine signaling networks, and HLA diversity, thereby providing a more predictive model for human vaccine response. Interestingly, the T cell response against PreS1 and PreS2 is much weaker in NHP than in CB6F1 mice, especially the CD4^+^ T cells response. Despite the low overall CD4^+^ T, and possibly Tfh response, the antibody response remained robust, and the memory B cell response was strong. These data suggests that LuS-specific Tfh cell may cross help the PreS1S2-specific B cells and hence compensate for the reduced level of PreS1S2-specific Tfh response, highlighting the advantage of the nanoparticle scaffolded antigen design.

Persistent antigenemia in chronic HBV infection is another well-established driver for T cell exhaustion, with high HBsAg level reported to be negatively correlated with response to therapeutic vaccination (*26, 67*). Therefore, immune-checkpoint blockade (ICB), an immune-stimulating approach widely employed in Oncology field, holds great potential as an adjunct to therapeutic vaccine. By overcoming immune-exhaustion, ICB may enhance efficacy of therapeutic vaccines, especially in patients with high level of HBsAg, and thus broaden the population of patients who can benefit from a therapeutic vaccine. Notably, in a recent clinical trial evaluating a combination of an siRNA (Imdusiran), a therapeutic vaccine (VTP-300), and an ICB (Nivolumab) demonstrated that the three-component regimen led to greater reduction of HBsAg level compared to the two-component regimen of siRNA and therapeutic vaccine. Given that mRNA-1965 is designed to elicit robust T cell response and promote function cure through cell-mediate mechanisms, both RNAi and ICB are compelling co-therapy to enhance its therapeutic efficacy. Indeed, our preliminary data has demonstrated that combining mRNA-1965 with an immune-stimulatory antibody cocktail, containing a PD-L1 antagonist and a OX40 agonist, significantly improved therapeutic outcomes when compared to mRNA-1965 monotherapy in mice with high baseline level of HBsAg. These findings support the use of ICB as a promising co-therapy for therapeutic vaccines for the treatment of chronic HBV infection.

We also observed that mRNA-1965 elicited markedly different immune profiles between mouse strains, with CB6F1 mice showing strongest CD8^+^ T cell response against PreS1S2, whereas C57BL/6 mice mounted strongest CD8^+^ T cell response against Core. Our findings highlight the stark difference in the landscape of cellular immune response to mRNA-1965 in CB6F1 and C57BL/6 strains of mice, likely driven by differences in MHC-restricted dominant epitopes and immune-hierarchy between different antigens. Such strain-dependent variability also underscores an inherent limitation of murine models, as differences in MHC haplotypes and immune repertoire restrict the extrapolation of mouse immune response to the human setting.

Furthermore, while mRNA-1965 only elicited minimal CD8+ T cell response to Polymerase in HBV-naïve C57BL/6 mice, a moderate CD8+ T cell response to Polymerase was observed in AAV mice of C57BL/6 background that eliminated HBV biomarkers post mRNA-1965 treatment. A previous study has similarly reported shifts in T cell response to different epitopes of Polymerase between naïve mouse and the AAV-HBV mouse (*68*). These findings suggest that the immune-landscape induced by mRNA-1965 is altered in AAV-HBV mice as compared to naïve mouse, potentially due to differential immune suppression among HBV antigens and epitopes.

It is well-established that achieving functional cure for CHB will likely require a multi-modal approach targeting distinct mechanisms, including viral replication inhibition, HBsAg reduction, and immune stimulation. In this paper, we demonstrate that combining mRNA-1965 and an antibody cocktail containing an OX40 agonist IgG and a PD-L1 antagonist IgG led to significantly enhanced therapeutic efficacy in AAV-HBV mice with high baseline antigenemia and viremia

Our study has several limitations. We observed interference, i.e. the immune response to one antigen blunted or diminished by the concurrent presence of another antigen, during the development of mRNA-1965. T cell responses against Core and Polymerase were substantially lower compared to responses elicited by monovalent mRNA vaccines administered at the same dose for the corresponding antigen. However, we were able to restore T cell responses to levels comparable to those induced by monovalent vaccines by increasing the total dose of mRNA-LNP while maintaining the relative ratio of each individual mRNA component. Albeit this data was only collected in naïve mice, it suggests that repeat immunization or higher dose level may be necessary for in vivo efficacy. The mRNA platform is particularly well-suited for repeat dosing because it does not induce anti-vector immunity, unlike viral-vectored vaccines that require heterologous prime-boost strategies (*69, 70*). Our NHP study has a small sample size, which consequently restricted the extent to which human HLA diversity is represented in the study. Nevertheless, this study is complementary to our in-silico analysis of the breath of cellular immune-response elicited by mRNA-1965 in human and effectively serves as a confirmatory study supporting the broad immunogenicity profile of mRNA-1965. Finally, our in-vivo therapeutic efficacy study has a relatively short duration, as the study was concluded three weeks after the final dose of mRNA-1965. A longer-term study is warranted to enable longitudinal monitoring of serum HBV biomarkers following sero-clearance. Nevertheless, the complete elimination of HBV-core positive hepatocytes in AAV-HBV mice that reached sero-clearance provides strong evidence of functional cure, while the marked reduction of HBV-core positive hepatocytes in high-antigenemia AAV-HBV mice treated with the antibody cocktail suggested the potential for sustained HBsAg reduction. Regarding the immune checkpoint blockade study, one limitation is the absence of treatment arms evaluating mRNA-1965 with each individual antibody, which would allow a more granular assessment of the contribution of each immune-stimulation; these studies are currently on-going. The on-going and additional studies will provide critical insights on how mRNA-LNP therapeutic vaccines synergize with other modalities to overcome immune-dysfunction and drive functional cure in chronic Hepatitis B.

In conclusion, the mRNA technology offers a significant advantage for the treatment of chronic HBV, as it can be adapted by encoding designer antigens and by combining it with additional immunotherapies to elicit the key signatures required for immune clearance. mRNA-1965 has the potential to steer the human immune system towards functional immunity, an aspect that needs to be evaluated in human clinical studies.

## Materials and Methods

### Vaccine design and synthesis

mRNAs encoding Surface, Core, and Polymerase were designed from HBV genotype A serotype ayw (NCBI accession no. LC592168), genotype D (NCBI accession no. OK106256), and genotype D (NCBI accession no. MW999652), respectively. Briefly, the mRNAs were formulated in a mixture of four lipids: an ionizable lipid, a neutral lipid, a sterol, and a PEG-modified lipid, in buffered sucrose. The formulated mRNA-lipid nanoparticles (LNPs) were stored at −60° to −90°C until use.

### Ethics, animals, treatments (includes AAV-HBV model)

Female CB6F1 mice, 8-week-old, were obtained from Charles River Laboratories, and used for all mouse studies except ones with AAV-HBV mouse model. All animal experiments were approved and conducted in accordance with the regulations of the Sponsor’s IACUC Protocol 18-03-04. Immunization was performed by i.m. injection of right quadricep in a volume of 50 µL.

Male Cambodian-origin Cynomolgus macaques, 4-6-years-old years of age, were sourced from Bioqual, Inc. The study was performed by BIOQUAL, Inc. located at 12301 Parklawn Drive, Rockville, MD 20852 (BPD) (vivarium for acclimation and vaccination) and at 9600 Medical Center Drive, Suite 101, Rockville, MD 20850(MCD) (in-vitro BSL-2 laboratories), all animal procedures were approved and conducted in accordance with the regulations of the contract research organization’s IACUC Protocol 24-010. Immunization was performed by i.m. injection of right quadricep according to contract research organization’s SOP NP-014.

The AAV-HBV mouse model study was performed by Creative Biolabs. Male C57BL6/J mice, 5-6-weeks of age were sourced from Charles River, Beijing, and CD45.1 mice sourced from Shanghai Model Organisms Center, Inc. The procedures that were applied to animals in this protocol were approved by CBL IACUC. The care and use of animals were conducted in accordance with the regulations of the AAALAC. All animal procedures were approved and conducted in accordance with the regulations of the contract research organization’s IACUC Protocol VST-SY-12. Immunization was performed by i.m. injection of right quadricep according to the contract research organization’s SOP VS-PPD-SOP-007.

### AAV-HBV model

Male C57BL/6J mice, 5-6-week-old, were obtained from Charles River, Beijing. All animal experiments were approved and conducted in accordance with the regulations of the contract research organization’s IACUC Protocol VST-SY-12. On Day -30, all animals received a dose of 5E8 vg, 1E9 vg, 5E9 vg, or 1E10 vg of viral vectors of recombinant AAV-HBV (rAAV8-1.3HBV, genotype D, BrainVTA, Cat No. PT-0717) via intravenous injection. On Day 0, all animals received an initial dose of 15 ug of mRNA-1965 via intramuscular injection of right quadricep in a volume of 50 µL. Control animals were injected with PBS. Three repeat doses were administered on Days 14, 28, and 42. Serum samples were collected from all animals on days -30, -5, 14, 28, 42, 63. Spleens were collected from all animals per group on Day 63 and immediately processed.

### HBV neutralization assay

HepG2 cells stably expressing the NTCP receptor are cultured in NTCP Media (DMEM with 10% FBS and 8 µg/mL Puromycin) overnight in clear, collagen-I coated 96-well plates (1.5e4 cells/well). Cells are then conditioned with Pre-infection Media (DMEM with 3% FBS, 2% DMSO, and 1X NEAA) and incubated overnight prior to viral infection with live HBV-D (ayw). On the day of viral infection, serum dilutions are incubated with ∼800 genome equivalents (GEQ) per well of HBV in Infection Media (DMEM with 3% FBS, 2% DMSO, 1X NEAA, and 4% PEG-8000) for 1 hour at RT. HepG2 cells are inoculated with the serum/virus mixture and incubated overnight. Cells are then washed 3 times with PBS and maintained in Infection Media for an additional 5 days. On day 6 post-infection, the cell supernatant is heat-inactivated at 56.0°C for 30 minutes and then used to assess secreted Hepatitis B e antigen (HBeAg) levels. HBeAg levels are normalized to the virus infection control and IC50 (NT50) is determined using GraphPad Prism.

### Anti-PreS1S2 IgG ELISA

96-well Nunc Maxisorp ELISA plates (Thermo Fisher, Cat#439454) were coated with 100ng/well of PreS1S2 (genotype D) protein (Moderna, Lot#2024/3.8). After overnight incubation at 4°C, plates were washed four times with 1x PBS/0.05% Tween-20 (Boston Bioproducts, Cat#IBB-171) and blocked for 1.5 hour at 37°C using SuperBlock (Thermo Fisher, Cat# 37515). After washing, five-fold serial dilutions of mouse serum were added to plate in 1x assay diluent (1xPBST+ 5% goat serum). Plates were incubated for 2 hours at 37°C, washed and horseradish peroxidase-conjugated goat anti-mouse immunoglobulin G (IgG) (Southern Biotech, Cat# 1030-05) was added at a 1:10,000 dilution in assay diluent. Plates were incubated for 1 hour at 37°C, washed, and bound antibody was detected with a 3,3’,5,5’-tetramethylbenzidine (TMB) substrate (SeraCare, Cat#5120-0077). After incubation for 10 minutes at room temperature, the reaction was stopped by adding a TMB stop solution (SeraCare, Cat# 5150-0021) and the absorbance was measured at 450nm. Titers were determined using a four-parameter logistic curve fit in Prism V10.4 (GraphPad 112 Software, Inc) and defined as the reciprocal dilution at approximately optical density 450 = 1 (normalized to a mouse standard on each plate).

### Intracellular Cytokine Staining (ICS)

Fresh or cryopreserved splenocytes were stimulated with peptide pools (2 ug/mL), mitogens or DMSO in the presence of protein transport inhibitors for 6 hours. Surface and intracellular staining were performed using a panel of fluorescently labeled antibodies targeting various T cell markers and cytokines. Functional markers included CD107a, IFN-γ, TNF-a, IL-2, IL-4, IL-5, IL-13, and CD44. Data acquisition was performed on Cytek Aurora flow cytometers, and spectral unmixing was conducted using Spectroflo software. Gating and Boolean analysis were performed using OMIQ.

### Assessment of antigen-specific memory B cells by flow cytometry

Cryopreserved peripheral blood mononuclear cell (PBMC) samples collected on Days - 43, 14, 42, and 70 were thawed and stained with monoclonal antibodies against CD3, CD8, CD14, CD56, CD19, CD20, CD21, CD27 and CD38. Recombinant protein of PreS1S2 from HBV genotype D was biotinylated and conjugated with either PE or APC-streptavidin, according to the manufacturer’s instruction (BioLegend), to form tetramers to bait for antigen-specific B cells. Non-B cells and dead cells were gated out with CD3, CD8, CD14, CD56 and LIVE/DEAD staining. Memory B cells were gated on as CD19+, CD20+ and then further differentiated into CD27+CD21+ and CD27+CD21lo populations.

### Immunochemistry, Image Acquisition, and Image analysis

IHC, conducted to visualize Hepatitis B core protein expression in liver tissues, was performed on formalin-fixed paraffin embedded (FFPE) sections using the Leica Bond RX autostainer (Leica Microsystems, Buffalo Grove, IL). Sections were baked and deparaffinized on the instrument, followed by an epitope retrieval for 20 minutes using Leica Epitope Retrieval Buffer 1 (cat no. AR9961). Dako serum free protein block (cat no. X090930-2, Agilent Dako, Santa Clara, CA) was incubated on the slides for 15 minutes at room temperature. Anti-HBcAg antibody (cat no. LS-C312204, LifeSpan Biosciences, Lynnwood, WA) was used at 1.0 ug/mL dilution at room temperature for 30 minutes. Secondary antibody and detection were performed using the Bond Polymer Refine Detection Kit (cat no. DS9800, Leica Microsystems, Buffalo Grove, IL). Bond DAB Enhancer (cat no. AR9432, Leica Microsystems, Buffalo Grove, IL) and bluing reagent (cat no. 3802918, Leica Microsystems, Buffalo Grove, IL) were used to enhance the color.

Whole-tissue slides were scanned at 20X magnification with the Pannoramic 250 Flash III (3DHISTECH, Budapest, Hungary) digital slide scanner. Digital images were analyzed for Hepatitis B core quantification using Halo (Indica Labs) software. First, manual annotations were created to identify individual liver regions per slides and next to detect positive HBcAg cells, the HALO Multiplex IHC v3.4.9 module was optimized to detect nuclear stain and anti-HBcAg immunolabeling. The results are expressed as the percentage (%) of HBcAg cell positivity = HBcAg positive cells/ total number of cells.

### T cell exhaustion phenotyping by flow cytometry

Cryopreserved splenocytes were thawed in complete media, washed with PBS, and stained with LIVE/DEAD Fixable UV Blue Dead Cell Stain for 15 minutes at room temperature. Surface staining was performed by incubating cells with an antibody cocktail prepared in Brilliant Stain Buffer, containing FC Block, CD44 BUV395, CD3 BUV737, CD8 BUV805, PD-1 BV421, CD4 BV480, CD69 BV711, I-A/I-E AF488, LAG-3 RB705, TIM-3 RB780, CTLA-4 PE-CF594, CD45.1 PE-Cy7, and SLAMF6 APC for 15 minutes at room temperature. Cells were washed twice in FC buffer, resuspended in 0.5% PFA-FC stain buffer, and filtered through a 30 µm filter in a 96-well plate before acquisition. Samples were acquired on an Aurora Spectral Flow Cytometer (Cytek Biosciences).

### Meso Scale Discovery (MSD) cytokine assay

Fresh splenocytes were stimulated with overlapping HBV peptide pools (2 ug/mL), mitogens, or DMSO for 6 hours. Cell culture supernatants were collected for all stimulation conditions and analyzed with a customized 8-plex MSD U-plex assay, including IFN-g, TNF-a, IL-2, IL-4, IL-5, IL-13, and IL-17a per manufacturer protocol.

### Pentamer staining

Fresh splenocytes were washed with PBS and stained with LIVE/DEAD Fixable UV Blue Dead Cell Stain for 15 minutes at room temperature. Following a wash with FC stain buffer, Core93-100 PE pentamer (Core93-100 Pentamer (MGLKFRQL) ProImmune (1431)) staining was performed in the presence of FC Block for 15 minutes at room temperature. After centrifugation, surface staining was performed by incubating cells with an antibody cocktail prepared in Brilliant Stain Buffer, containing FC Block, CD44 BUV395, CD3 BUV737, CD8 BUV805, PD-1 BV421, CD4 BV480, BV711, I-A/I-E AF488. Cells were fixed in 0.5% PFA and acquired on an Aurora Spectral Flow Cytometer (Cytek Biosciences).

### Quantification of HBsAg, HBeAg, HBV DNA, and anti-HBs IgG titer

Serum levels of HBsAg, HBeAg, HBV DNA, and anti-HBs IgG were measure using respective kits per manufacturer protocol by Creative Biolabs: HBeAg Kit (Maccura, Cat No. IM4403003); HBsAg kit (Maccura, Cat No. IM4403001); Anti-HBs (HBsAb) kit (Maccura, Cat No. IM4403002); Hepatitis B Virus Nucleic Acid Kit (Sansure, Cat No. 20153400083).

### In vivo killing assay

Splenocytes were harvested from donor CB6F1 mice and pulsed with 10 µg/mL of pre-identified HBV-specific epitopes. Pulsed cells were mosaically labeled with CellTrace Violet (CTV+/++) or CellTrace Far Red (CTFR+) to distinguish specific populations, and co-stained with CD45.2 PE-Cy7 for identification. 2e7 labeled peptide-pulsed splenocytes were intravenously transferred into vaccinated recipient CB6F1 mice on Day 29. Sixteen hours post-transfer (Day 30), spleens were harvested from recipient CB6F1 mice and analyzed by flow cytometry. Panel for flow cytometry includes LIVE/DEAD Fixable UV Blue, CD3 BV605, CD4 BV480, CD8 BUV805, I-A/I-E AF488, and CD45.2 PE-Cy7. Data were acquired on a Cytek Aurora Spectral Flow Cytometer. Killing was calculated as:

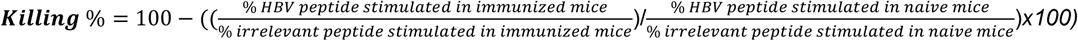

where target cell ratios were determined from CTV/CTFR-labeled populations. The approach enabled precise quantification of antigen-specific killing mediated by HBV-specific T cells.

### Statistical analyses

A two-way ANOVA with Tukey’s multiple comparison correction was used to test for differences in T cell responses against HBV antigens of different genotypes. A one sample t-test with a Benjamini-Hochberg multiple test correction was used to test for differences in HBV biomarkers in serum of AAV-HBV mice.

## Acknowledgements

D.M., Y.S., A.DiP., and S.P. conceptualized and supervised the research; D.M., Y.S., H.D., L.M., T.K., E.D., M.M., E.O., M.C., C.A., P.M., C.H., Z.W., G.A.A., A.F., L.M., and Y.L. performed the research. K.H., G.A., A.C., reviewed and validated the research. D.M., Y.S., A.DiP, and S.P. wrote the manuscript and designed the figures. All authors reviewed the manuscript.

All flow cytometric data were generated in the Flow Cytometry Shared Resource Laboratory, managed by the Flow Cytometry & Immunoassay Group within the Research Automation team at Moderna Therapeutics.

