## Supplementary for "A multivalent mRNA-LNP therapeutic vaccine with broad cross-genotype immunogenicity elicits clearance of HBV infected hepatocytes"

### Supplementary Materials and Methods

#### Antibody-dependent complement deposition (ADCD)

To perform Antibody-dependent complement deposition (ADCD), antigen-coupled beads were incubated with mouse serum for 2 h at 37°C to form immune complexes. The immune complexes were washed, and lyophilized guinea pig complement (Cedarlane) in gelatin veronal buffer with calcium and magnesium (GBV++) (Boston BioProducts) was added for 30 min (complement was reconstituted according to manufacturer's instruction). The deposition of complement was detected by fluorescein-conjugated goat IgG fraction to guinea pig Complement C3 (Mpbio). ADCD was reported as the median of C3 deposition.

#### Antibody-dependent cellular phagocytosis (ADCP)

Antibody-dependent cellular phagocytosis was performed as previously described with slight modifications. Briefly, antigens (PreS1S2 and S-HBsAg) were coupled to yellow-green beads (Thermofisher). Serum samples were diluted in culture medium and incubated with antigen-coated beads for 2 h at 37°C to form immune complexes. THP-1 (ATCC) cells (25,000/well) were added to the immune complexes and incubated at 37C for 4 h. Cells were washed 2x with cold 1x PBS prior to being fixed with 4% paraformaldehyde. The cells were analyzed on iQue (Intellicyt). Analysis was performed on IntelliCyt ForeCyt (v8.1). PE median fluorescent intensity (MFI) is reported as a readout for antigen-specific antibody. Buffer only wells were used as negative controls and the assay was performed in technical replicate with two biological replicates

#### Epitope Mapping

CB6F1 mice were immunized twice with mRNA-LNP encoding for either HBV Large (A), Core (D) or Polymerase (D). On Day 36, spleens were harvested and Epitope mapping of the PreS1S2, Core, and Polymerase antigens was performed by stimulating cells with individual 15- AA peptides that overlapped by 11 AA to covering the entire consensus wild-type proteins. T cell ICS was performed using the methods previously stated to down select on immunogenic CD8 IFN-y+ peptides hits.

#### Human epitope prediction

CD8$^+$ T cell epitope predictions were performed for Hepatitis B virus antigens (PreS1, PreS2, Small, Core_149, and Polymerase_8mut) across genotypes A–D using NetMHCpan 4.1 to evaluate peptide–HLA binding affinity (*71*). Predicted epitopes were assessed against a representative panel of HLA class I alleles covering the global population, and strong-binding epitopes were retained. Population coverage was estimated based on allele frequency data from the Allele Frequency Net Database (AFND) (*72*), where each peptide's restricting HLA alleles were weighted by their observed frequencies to calculate cumulative global coverage. Epitopes predicted to achieve ≥80% population coverage were mapped along each protein sequence to identify regions of high epitope density across HBV genotypes.

#### Serum ALT level measurement

Serum ALT level was measured using Alanine Transaminase Activity Assay Kit (Abcam, ab105134) per manufacturer protocol.

#### Statistical analyses

A two-way ANOVA with Tukey's multiple comparison correction was used to test for differences in antibody binding titers against PreS1S2 and S-HBsAg; and in T cell responses against HBV antigens of different genotypes. A one sample t-test with a Benjamini-Hochberg multiple test correction was used to test for differences in HBV biomarkers in serum of AAV-HBV mice.

**Figure S1**

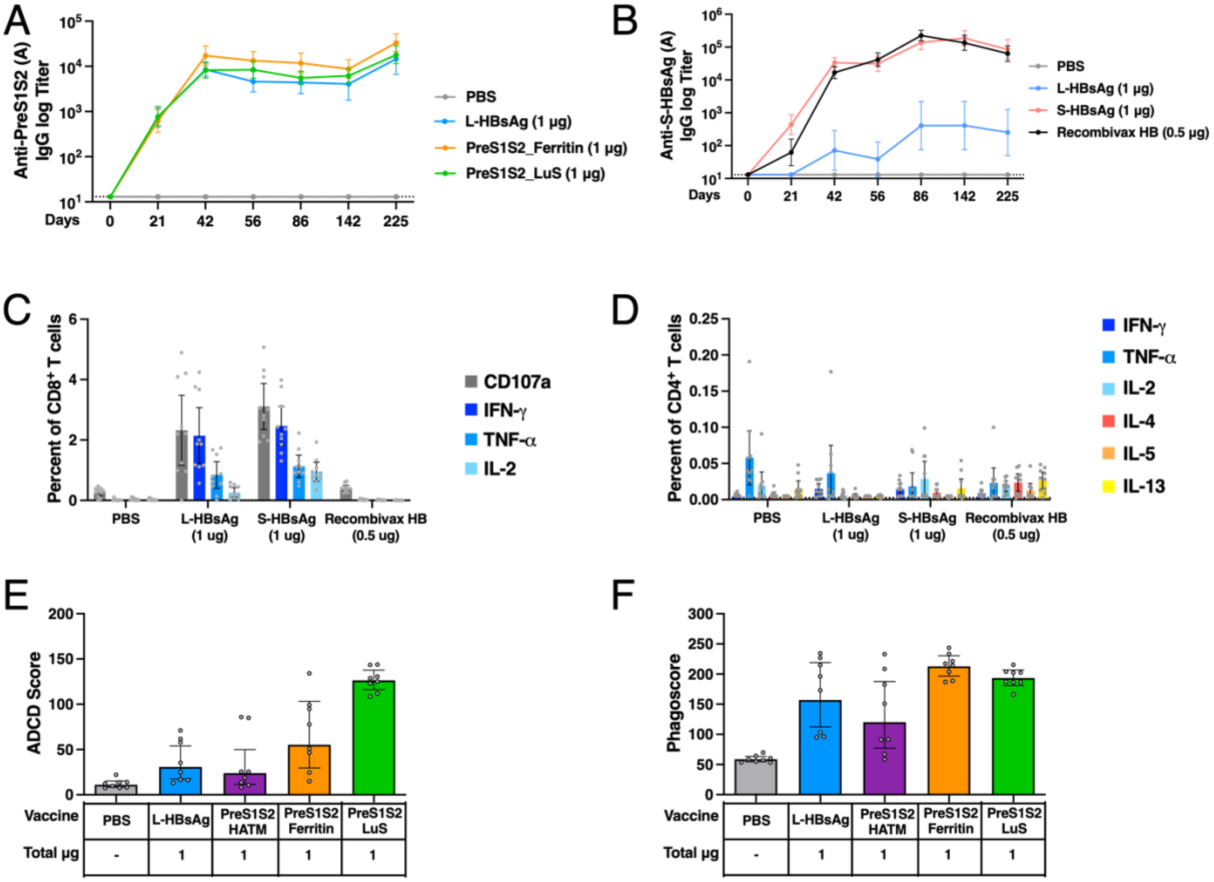

A
Anti-PreS1S2 (A) IgG log Titer
10^5
10^4
10^3
10^2
10^1
Days
0
21
42
56
86
142
225
PBS
L-HBsAg (1 µg)
PreS1S2_Ferritin (1 µg)
PreS1S2_LuS (1 µg)
B
Anti-S-HBsAg (A) IgG log Titer
10^6
10^5
10^4
10^3
10^2
10^1
Days
0
21
42
56
86
142
225
PBS
L-HBsAg (1 µg)
S-HBsAg (1 µg)
Recombivax HB (0.5 µg)
C
Percent of CD8+ T cells
6
4
2
0
PBS
L-HBsAg (1 ug)
S-HBsAg (1 ug)
Recombivax HB (0.5 ug)
CD107a
IFN-γ
TNF-α
IL-2
D
Percent of CD4+ T cells
0.25
0.20
0.15
0.10
0.05
0.00
PBS
L-HBsAg (1 ug)
S-HBsAg (1 ug)
Recombivax HB (0.5 ug)
IFN-γ
TNF-α
IL-2
IL-4
IL-5
IL-13
E
ADCD Score
200
150
100
50
0
Vaccine
PBS
L-HBsAg
PreS1S2 HATM
PreS1S2 Ferritin
PreS1S2 LuS
Total µg
-
1
1
1
1
F
Phagoscore
300
250
200
150
100
50
0
Vaccine
PBS
L-HBsAg
PreS1S2 HATM
PreS1S2 Ferritin
PreS1S2 LuS
Total µg
-
1
1
1
1

**Figure S1. Additional immunogenicity data on HBsAg antigen constructs.**

Female CB6F1 mice received two intramuscular 1 ug doses of HBV mRNA-LNP constructs or 0.5 ug doses of Recombivax or PBS on days 0 and 21 as described in Fig 1. (**A-B**) Anti-PreS1S2 (**A**) and S-HBs (**B**) HBV genotype A IgG binding titer was measured using serum samples collected from days 21, 42, 56, 86, 142, and 225. (**B**) Neutralization titer was measured using day 225 serum with or without prior incubation with 33ug/mL of S-HBs SVP. (**C-D**) CD8 and CD4 T cell response against S-HBs peptides of HBV genotype A was measured by ICS. (**E**) Antibody-dependent compliment deposition (ADCD) score. (**F**) Antibody-dependent cellular phagocytosis (ADCP) measured using day 42 serum. Plots in panels A,B,E,F show group geometric mean with 95% CI, n=10 per group. Plots in panels C-D show group mean with 95% CI, n=10 per group. Assay LLOD indicated by dashed lines.

### Figure S2

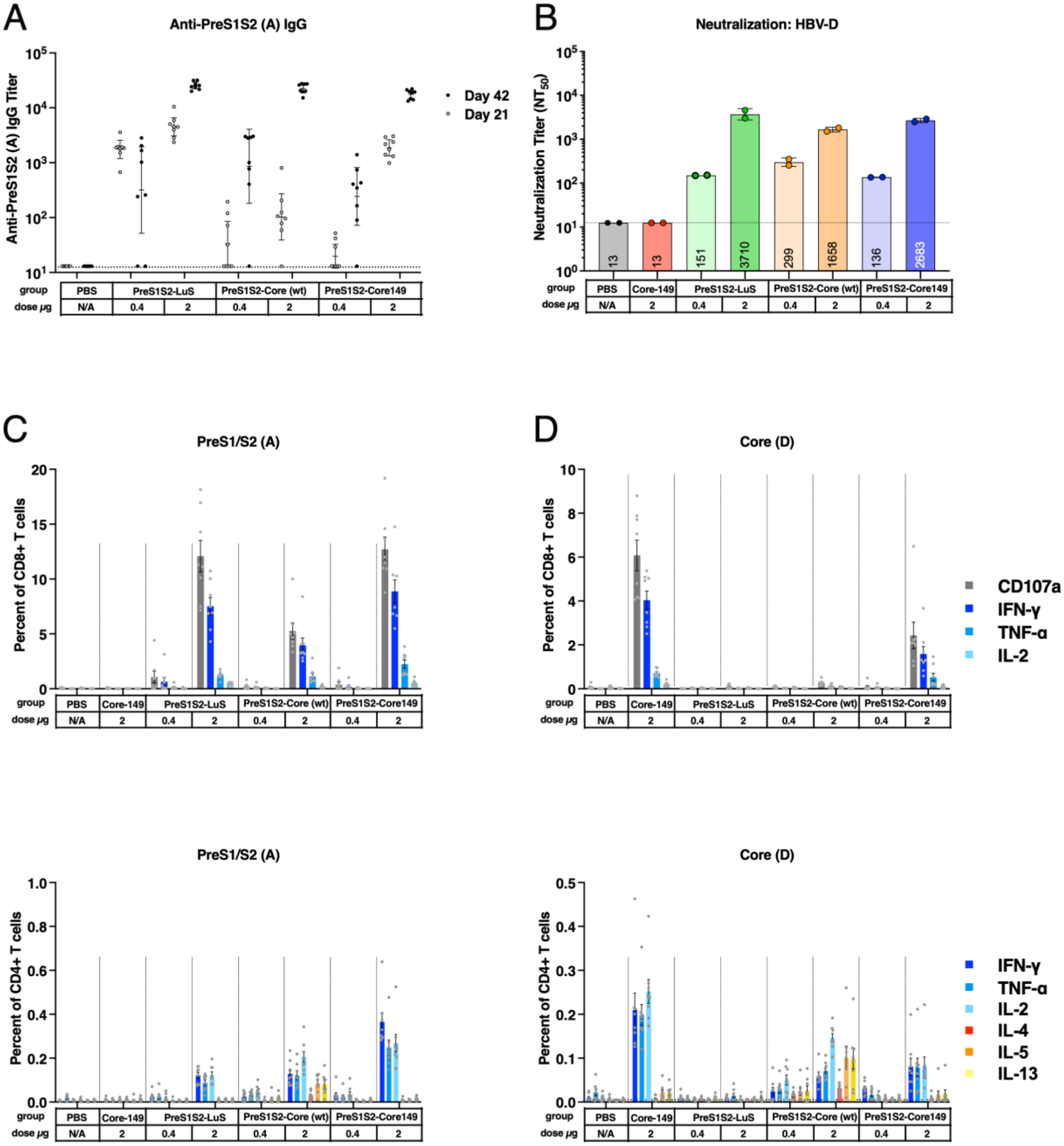
A
Anti-PreS1S2 (A) IgG
Anti-PreS1S2 (A) IgG Titer
Day 42
Day 21
B
Neutralization: HBV-D
Neutralization Titer (NT50)
13
13
151
3710
299
1658
136
2683
C
PreS1/S2 (A)
Percent of CD8+ T cells
D
Core (D)
CD107a
IFN-γ
TNF-α
IL-2
PreS1/S2 (A)
Percent of CD4+ T cells
Core (D)
IFN-γ
TNF-α
IL-2
IL-4
IL-5
IL-13
group
dose μg
PBS
Core-149
PreS1S2-LuS
PreS1S2-Core (wt)
PreS1S2-Core149
N/A
0.4
2

**Figure S2. Immunogenicity of PreS1S2-Core chimeric constructs**

Female CB6F1 mice received two intramuscular doses of HBV mRNA-LNP constructs or PBS on days 0 and 28. (**A**) Anti-PreS1S2 binding antibodies at days 21 and 42. (**B**) HBV genotype D neutralization titer measured using day 42 serum. (**C-D**) CD8 and CD4 T cell response against PreS1S2 peptides of HBV genotype A and Core peptides of HBV genotype D, measured by ICS. Plots in panels A show group geometric mean with 95% CI, n=10 per group. Plots in panel B show mean of two replicates, n=10 per group. Plots in panels C-D show group mean with 95% CI, n=10 per group. Assay LLOD indicated by dashed lines.

**Figure S3**

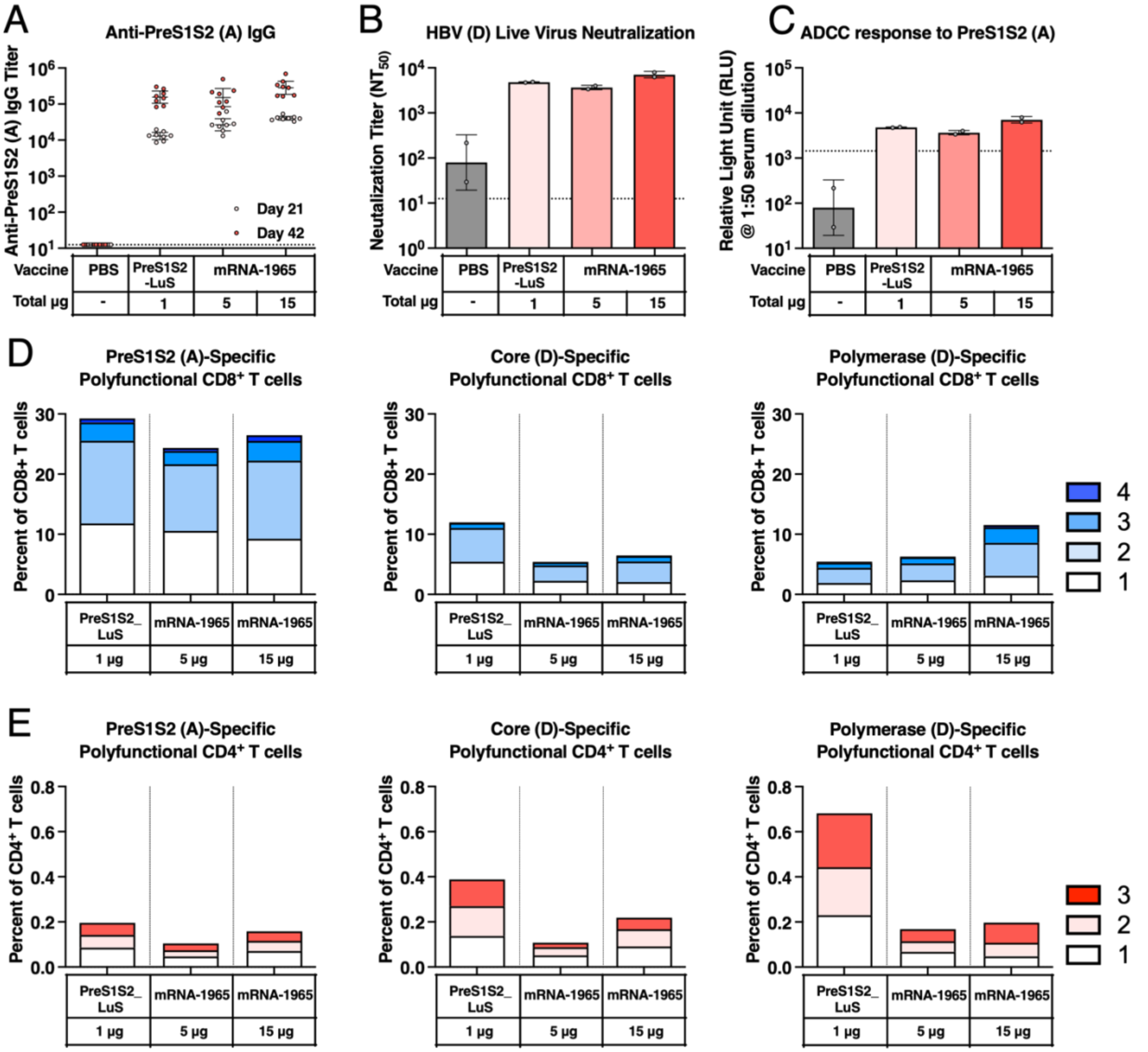
A
Anti-PreS1S2 (A) IgG
Anti-PreS1S2 (A) IgG Titer
Day 21
Day 42
Vaccine
PBS
PreS1S2 -LuS
mRNA-1965
Total µg
-
1
5
15
B
HBV (D) Live Virus Neutralization
Neutalization Titer (NT50)
C
ADCC response to PreS1S2 (A)
Relative Light Unit (RLU) @ 1:50 serum dilution
D
PreS1S2 (A)-Specific Polyfunctional CD8+ T cells
Core (D)-Specific Polyfunctional CD8+ T cells
Polymerase (D)-Specific Polyfunctional CD8+ T cells
Percent of CD8+ T cells
PreS1S2_ LuS
1 µg
5 µg
15 µg
4
3
2
1
E
PreS1S2 (A)-Specific Polyfunctional CD4+ T cells
Core (D)-Specific Polyfunctional CD4+ T cells
Polymerase (D)-Specific Polyfunctional CD4+ T cells
Percent of CD4+ T cells

**Figure S3. mRNA-1965 display immunogenicity comparable to monovalent mRNA vaccines.**

Female CB6F1 mice received two intramuscular doses of mRNA-1965 or respective monovalent mRNA constructs of HBV antigens at indicated doses or PBS on days 0 and 28. (**A**) Anti-PreSeS2 HBV genotype A IgG binding titer was measured using serum samples collected from days 21 and 42. (**B-C**) Serum neutralization titer (**B**) and ADCC induction (**C**) was measured using serum samples collected from day 42. (**D-E**) Antigen-specific CD8+ (**Top**) and CD4+ (**Bottom**) T cell response against PreS1S2 of HBV genotype A (**Left**), Core (**Middle**) and Polymerase (**Right**) of HBV genotype D using splenocyte samples collected on Day 42. Colors within each bar indicates the proportion of antigen specific T cells capable of producing respective number of CD8+ T cell marker/cytokine (CD107a, IFN-γ, TNF-α, IL-2) or Th1 CD4+ T cell cytokines (IFN-γ, TNF-α, IL-2).

**Figure S4**

A

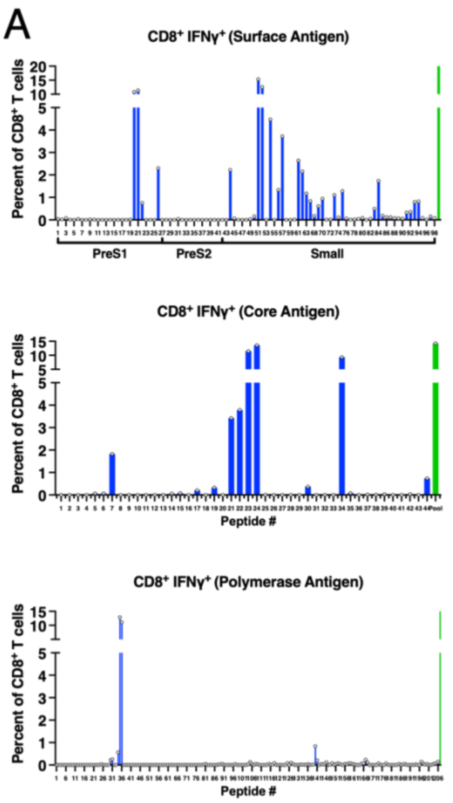


B

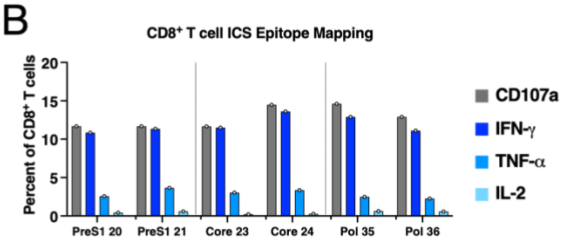


C

| Peptide | CD8 IFN-y+ (%) | AA Sequence | AA Start/Stop | Specificity |
|---|---|---|---|---|
| 20 | 10.9 | WSPQAQGILTTVSTI | 77-91 | PreS1 |
| 21 | 11.4 | AQGILTTVSTIPPPA | 81-95 | PreS1 |
| 23 | 11.5 | VNTNMGLKFRQLLWF | 89-103 | Core |
| 24 | 13.6 | MGLKFRQLLWFHISC | 93-107 | Core |
| 35 | 12.9 | LVNHYFQTRHYLHTL | 137-151 | Pol |
| 36 | 11.1 | YFQTRHYLHTLWKAG | 141-155 | Pol |

D

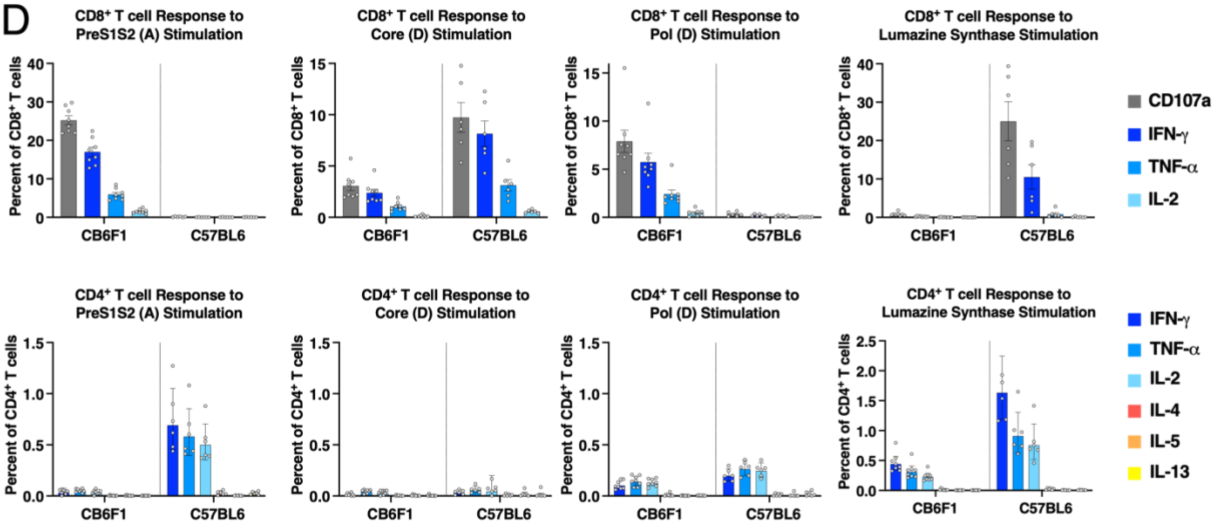

**Figure S4**. **Epitope mapping in CB6F1 mice and Differences in Immunogenicity between C57BL/6 and CB6F1 mouse strains.**

(**A-C**) Mice received two doses of monovalent mRNA constructs, encoding Large-HBs, WT core, and WT Polymerase, at 10ug on days 0 and 28. Splenocytes were collected and pooled on day 42 for epitope mapping. (**A**) CD8+ T cell response against each epitope in HBV Surface (**Top**) (genotype A), Core (**Middle**) and Polymerase (**Bottom**) (genotype D) by IFN-g. Green bars indicate responses to peptide pools covering the entire antigen. (**B**) CD8+ T cell response against dominant epitopes in each antigen. Colors of bars indicate respective cytokines. (**C**) Table of AA sequences of the two most immunogenic 15-mer peptides of each antigen. (**D**) CB6F1 or C57BL/6 mice received two doses of mRNA-1965 (15ug) on days 0 and 28. CD8+ (**Top**) and CD4+ (**Bottom**) T cell response against PreS1S2 (**Left**) genotype A, Core (**2$^{nd}$ to Left**) and Polymerase (**2$^{nd}$ to Right)** (genotype D), and Lumazine Synthase (**Right**) were measured using splenocyte samples collected on Day 42. Bar heights and error margins represent geometric means and 99% CI. Colors of bars indicate respective cytokines. N=8 for CB6F1 group and n=6 for C57BL/6 group.

**Figure S5**

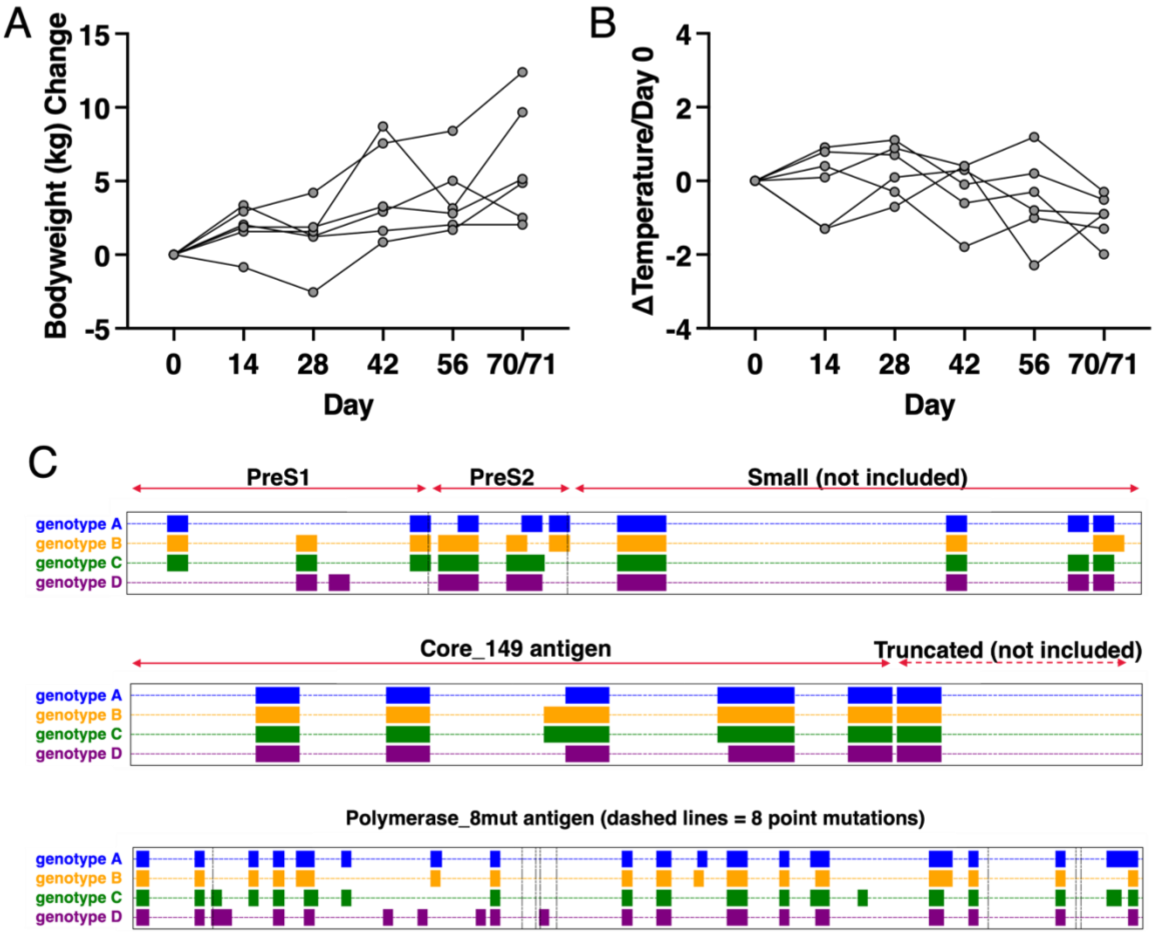

A
Bodyweight (kg) Change
15
10
5
0
-5
0
14
28
42
56
70/71
Day
B
ΔTemperature/Day 0
4
2
0
-2
-4
0
14
28
42
56
70/71
Day
C
PreS1
PreS2
Small (not included)
genotype A
genotype B
genotype C
genotype D
Core_149 antigen
Truncated (not included)
genotype A
genotype B
genotype C
genotype D
Polymerase_8mut antigen (dashed lines = 8 point mutations)
genotype A
genotype B
genotype C
genotype D

**Figure S5**. **Safety assessment of NHPs following mRNA-1965 immunization and high-coverage HBV epitope prediction.**

(**A-B**) Male Cynomolgus Macaques, 4-6 years old, received three intramuscular doses of mRNA-1965 at 150ug on days 0, 28, and 56. Changes in body weight (**A**) and temperature (**B**) of each NHP were recorded on days 0, 14, 28, 42, 56, and 70. Red arrows indicate time of vaccinations. (C) Epitopes in HBV Surface, Core, and Polymerase antigens that's predicted to cover over 80% world population based on pMHC binding affinity and global HLA frequencies.

**Figure S6**

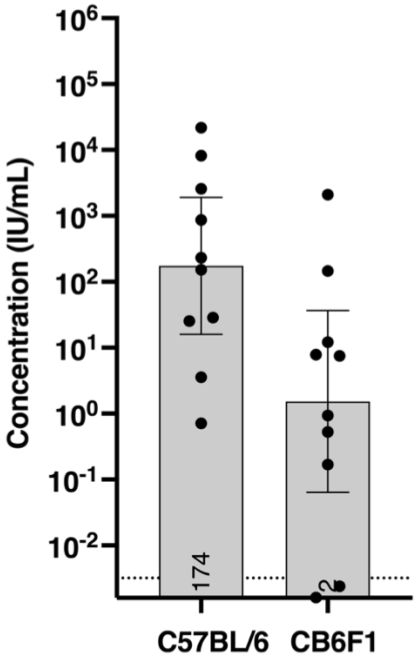

Concentration (IU/mL)
10^6
10^5
10^4
10^3
10^2
10^1
10^0
10^-1
10^-2
174
2
C57BL/6
CB6F1

**Figure S6**. **Evaluation of serum HBsAg in CB6F1 and C57B6 mice following AAV-HBV dosing.**

Male C57BL/6 or CB6F1 mice, 6-7 weeks old, were injected through the tail vein with 1e10, vector genomes of recombinant AAV-HBV on Day 0. Serum samples were collected on Day 25, and serum HBsAg level was evaluated.

142 **Figure S7**

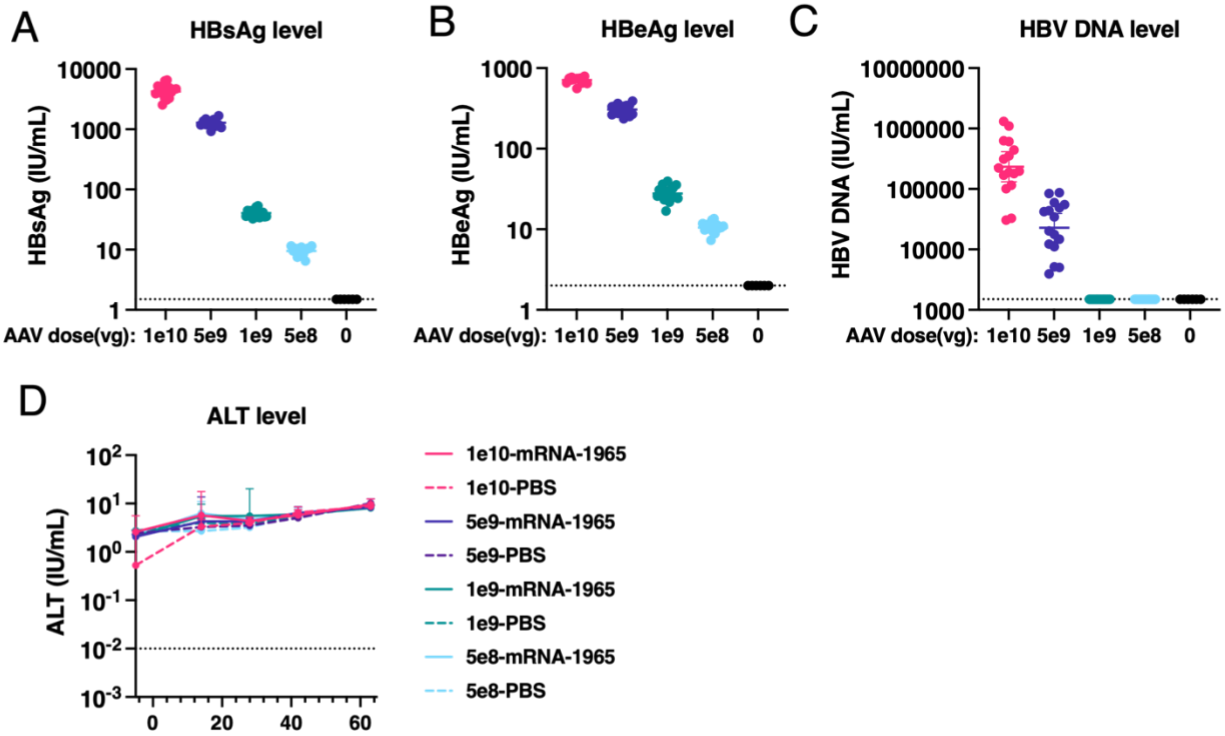


143

**Figure S7**. **AAV-HBV model characterization and ALT post treatment.**

Male C57BL/6 mice, 6-7 weeks old, were injected through the tail vein with 1e10, 5e9, 1e9, or 5e8 vector genomes of recombinant AAV-HBV on Day -30. Mice received four intramuscular doses of mRNA-1965 at 15ug or PBS on Days 0, 14, 28, and 42. Serum samples were collected on Days -5, 14, 28, 42, 63. (**A-C**) Serum HBsAg (**A**), HBeAg (**B**), and HBV DNA (**C**) levels were measured using Day -5 serum samples. Center bold lines and error margins represent group geometric means and 95% CI. (D) Serum ALT levels across time points. Dots and error margins represent group geometric means and 95% CI.
